# The catalytic-independent function of LSD1 modulates the epigenetic landscape of mouse embryonic stem cells

**DOI:** 10.1101/2023.06.29.547010

**Authors:** Sandhya Malla, Kanchan Kumari, Carlos Martinez-Gamero, Carlos A. García-Prieto, Stephanie Stransky, Jonatan Caroli, Damiana Álvarez-Errico, Paulina Avovome Saiki, Weiyi Lai, Cong Lyu, Jonathan D. Gilthorpe, Hailin Wang, Simone Sidoli, Andrea Mattevi, Andre Mateus, Manel Esteller, Angel Roman, Francesca Aguilo

## Abstract

Lysine-specific histone demethylase 1 (LSD1), which demethylates mono- or di-methylated histone H3 on lysine 4 (H3K4me1/2), is essential for early embryogenesis and development. Here we show that LSD1 is dispensable for embryonic stem cell (ESC) self-renewal but is required for ESC growth and differentiation. Reexpression of a catalytically-dead LSD1 (LSD1^MUT^) recovers the proliferation capability of ESCs, yet the enzymatic activity of LSD1 is essential to ensure proper differentiation. Indeed, a gain of H3K4me1 in *Lsd1* knockout (KO) ESCs does not lead to major changes in global gene expression programs related to stemness. However, ablation of LSD1 but not LSD1^MUT^ results in decreased DNMT1 and UHRF1 proteins coupled to global hypomethylation. We show that both LSD1 and LSD1^MUT^ control protein stability of UHRF1 and DNMT1 through interaction with the ubiquitin-specific peptidase 7 (USP7) and, consequently, inhibiting DNMT1 and UHRF1 ubiquitylation. Our studies elucidate for the first time a novel mechanism by which the scaffolding function of LSD1 controls DNA methylation in ESCs.

## Introduction

Embryonic stem cells (ESCs) represent the immortal *in vitro* capture of a very early stage of the developing embryo. They maintain the potential to proliferate indefinitely and differentiate into all three embryonic germ layers, providing material for cell-based therapies and holding out the promise to transform the next generation of medicine. In contrast to somatic cells, for which the transcriptional status of most genes is epigenetically fixed, ESCs globally possess decondensed chromatin, which can constantly be remodelled during developmental specification [1, 2]. This is sustained by the coordination of transcription factors, chromatin regulators, DNA and histone marks, and RNA modifiers [1, 3].

Lysine-specific histone demethylase 1 (LSD1; also known as KDM1A/AOF2/BHC110) is a histone modifying enzyme that demethylates the mono-and di-methyl moieties of histone H3 lysine 4 (H3K4me1/2) [4, 5]. Although LSD1 has been shown to be involved in early embryogenesis [6, 7], the function of this epigenetic factor in ESC self-renewal and differentiation is poorly understood [8]. For example, LSD1 has been shown to maintain ESC self-renewal by silencing developmental genes [9], and loss of LSD1 promotes neural lineage differentiation [10]. Conversely, it has been reported that mouse ESCs deleted of LSD1 retain stem cell characteristics, suggesting that LSD1 timely regulates the expression of key developmental regulators during early embryonic development [11]. In addition, other studies have also shown that LSD1 is not essential for the maintenance of ESC identity but it is required for the differentiation of multiple cell types *in vitro* and for the late cell-lineage determination and differentiation during pituitary organogenesis *in vivo* [7, 12–15]. Mechanistically, it has been proposed that LSD1 is poised at active pluripotency enhancers in ESCs to rapidly silence the pluripotency program during lineage commitment allowing for proper differentiation [16].

Non-histone substrates of LSD1, including key regulators of DNA methylation maintenance such as DNA (cytosine-5)-methyltransferase 1 (DNMT1) and Ubiquitin-like, with PHD and RING finger domains 1 (UHRF1), have also been demonstrated [6, 17–19]. Indeed, deletion of *Lsd1*/LSD1 in mouse ESCs and cancer cells have been shown to induce progressive global DNA hypomethylation [6, 20]. LSD1-mediated demethylation of DNMT1 increases DNMT1 protein stability by preventing its degradation by the proteasome [6, 21, 22]. Similarly, LSD1-mediated demethylation of UHRF1 in the G2/M phase prevents its ubiquitination and thus UHRF1 degradation [20], and diminishes the interaction between UHRF1 and PCNA, which is essential for DNA repair [17]. Yet another study proposes an indirect mechanism by which H3K4me1 demethylation by LSD1 is required to maintain DNA methylation levels at pluripotency genes leading to enhancer silencing during ESC differentiation [23].

Several lines of evidence demonstrate that demethylase-independent functions of LSD1 orchestrate tumorigenesis [24]. For instance, it has been shown that the interaction with LSD1 can lead either to degradation (e.g. p62) or stabilisation (e.g. ERRα) of the interacting protein independently of LSD1 catalytic activity [25]. In addition, LSD1 functions as a pseudosubstrate of the E3 ubiquitin ligase FBXW7 triggering its self-ubiquitination and rapid degradation [26]. More recently, the histone demethylase activity of LSD1 has been reported to be dispensable for endocrine-resistant breast tumorigenesis [27]. Whether LSD1 non-canonical mechanisms also operate in pluripotency is yet to be explored.

Here, we show that ablation of LSD1 in ESCs does not affect ESC self-renewal but impairs cellular growth. Albeit *Lsd1* KO cells with reexpression of a catalytically-dead LSD1 (LSD1^MUT^) recover the proliferation capability, both *Lsd1* KO and LSD1^MUT^ ESCs undergo defective differentiation. In ESCs, loss of LSD1 results in a gain of H3K4me1 in the promoters and distal intergenic regions of a subset of genes without affecting global gene expression programs related with stemness. Moreover, in the ESC state, deletion of *Lsd1* results in global hypomethylation coupled with decreased DNMT1 and UHRF1 protein levels. Notably, *Lsd1* KO with reexpression of wild-type LSD1 (LSD1^WT^) or LSD1^MUT^ recover the protein levels of DNMT1 and UHRF1 protein, and thus, DNA methylation levels. Strikingly, recovery of DNA methylation in LSD1^MUT^ is not sufficient to allow for normal differentiation. Mechanistically, LSD1^WT^ and LSD1^MUT^ can associate with the deubiquitinase USP7 (ubiquitin-specific protease 7, also known as HAUSP) and protect DNMT1 and UHRF1 from proteasomal degradation. Our results prompt a re-evaluation of the proposed mechanism of action for LSD1 in demethylating non-histone substrates, especially DNMT1 and UHRF1, to increase their stability. They also bring light to a new LSD1-USP7 axis to coordinate DNA methylation maintenance in mouse ESCs.

## Results

### *Lsd1* is dispensable for ESC self-renewal

To understand the regulatory function of LSD1 in pluripotency, we first assessed the expression of *Lsd1* in retinoic acid (RA)-induced neuronal differentiation and in embryoid bodies (EBs), comprising representatives of all three embryonic layers. Quantitative-reverse transcription PCR (RT-qPCR) revealed a moderate decrease in *Lsd1* mRNA during RA-induced differentiation whereas *Lsd1* mRNA expression was significantly higher at days 2, 4 and 6, but returned to control levels at day 8 of EB differentiation (**Figures 1A** and **1B**). LSD1 protein levels decreased at day 2 upon RA treatment but not during EB differentiation (**Figures 1C** and **1D**, upper panel), arguing for a specific post-transcriptionally regulation of LSD1 occurring during neural differentiation. OCT4 expression was used to monitor proper ESC differentiation (**Figures 1A-1D**). We next employed CRISPR/Cas9 to generate *Lsd1* knockout (KO) ESCs. To this aim, we used two different strategies: (i) a combination of two single guide RNAs (sgRNAs #1 and #2) targeting exon 1 (namely KO1); and (ii) one sgRNA (#3) targeting exon 6 which included the SWIRM domain of LSD1 (thereafter referred to as KO2; **Figure S1A**). After picking and expanding individual clones, targeted disruption of *Lsd1* was confirmed by PCR and Sanger sequencing (**Figures S1B** and **S1C**). Consistently, three bands were detected in KO1 whereas KO2 displayed two bands after T7 endonuclease I digestion, indicative of biallelic heterozygosity and homozygosity, respectively **(Figure S1D).** *Lsd1* KO mouse ESCs were further validated by western blotting and immunofluorescence analysis (**Figures S1E** and **S1F).**

**Figure 1.**
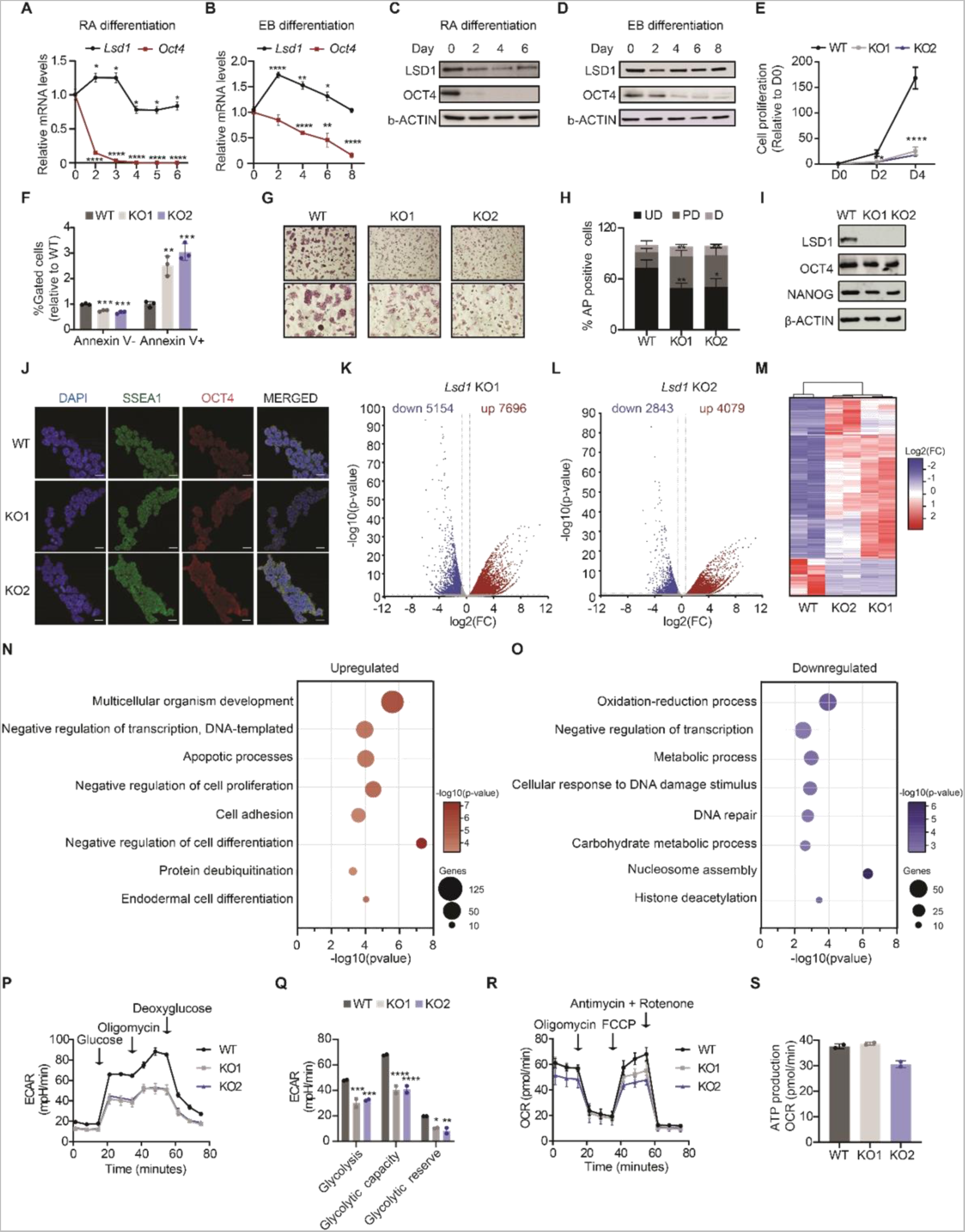
Loss of *Lsd1* does not lead to transcriptional deregulation of pluripotency genes. (A and B) RT-qPCR analysis of *Lsd1* and *Oct4* in mouse ESCs along the course of (A) retinoic acid (RA)-mediated and (B) embryoid body (EB) differentiation. mRNA levels are relative to the expression at day 0. (C and D) Western blot of LSD1 and OCT4 on the whole-cell extracts (WCE) of mouse ESC subjected to (C) RA or (D) EB differentiation. β-ACTIN is used as the loading control. (E) Relative cell proliferation rate of WT and *Lsd1* KO ESCs. (F) Percentage of live (Annexin V-) and apoptotic cells (Annexin V+) in *Lsd1* KO ESCs relative to WT. (G and H) (G) AP staining images and (H) quantification of colonies in WT and *Lsd1* KO ESCs. Undifferentiated (UD), partially differentiated (PD), and differentiated (D). Scale bars, 20μM. (I) Western blot of LSD1, OCT4, and NANOG on whole cell extracts of WT and *Lsd1* KO ESCs. β-ACTIN is used as the loading control. (J) Immunofluorescence images of SSEA1 and OCT4 in WT and *Lsd1* KO ESCs. DAPI was used as the nuclear marker. Scale bars, 20μM. (K and L) Volcano plots of differentially expressed genes in (K) *Lsd1* KO1 and (L) *Lsd1* KO2 ESCs in comparison to WT ESCs. Significant upregulated and downregulated transcripts are represented in red and blue, respectively (p < 0.05 and Fold change (FC) > 1.5). Non-significant hits are shown in grey dots. FDR value was calculated with the Benjamini–Hochberg correction. (M) Heatmap of differentially expressed genes in WT and *Lsd1* KO ESCs. The upregulated and downregulated genes are indicated in red and blue, respectively. (N and O) Gene ontology (GO) analysis of biological processes related to the common (N) upregulated and (O) downregulated genes of *Lsd1* KO ESCs compared to WT ESCs (p < 0.05 and FC > 1.5). (P and Q) (P) Measurement of extracellular acidification rate (ECAR) at indicated time points to determine glycolysis stress and (Q) glycolytic metabolic parameters in WT and *Lsd1* KO ESCs. (R and S) Quantification of Oxygen consumption rate (OCR) over time and (S) ATP production in *Lsd1* KO ESCs compared to WT ESCs using the Seahorse mito stress test. Statistical analysis: unpaired t-test (A-B and S), one-way ANOVA (E, F, H) and two-way ANOVA (Q). ∗p < 0.05, ∗∗p < 0.01, ∗∗∗p < 0.001, and ∗∗∗∗p < 0.0001. Error bars denote mean ± SD; n = 3 (A, B, E and F); n = 2 (P-S). Results are one representative of n = 3 independent experiments (C, D, G and J).

In order to characterise *Lsd1* KO clones, we performed proliferation, apoptosis, and cell cycle assays. Loss of LSD1 resulted in severely compromised cell growth and a marked increase in late apoptosis, suggesting that LSD1 is required for the normal proliferative function of ESCs (**Figures 1E** and **1F**). However, no significant difference in the cell cycle profile was observed in *Lsd1* KO compared to WT ESCs (**Figure S1G)**. In addition, *Lsd1* KO ESCs retained ESC morphology and were alkaline phosphatase (AP)-positive, indicative of maintenance of pluripotency (**Figure 1G**). Of note, *Lsd1* deletion increased the number of partially differentiated colonies, whereas the number of undifferentiated colonies significantly decreased (**Figure 1H**). Such an increase in the percentage of partially differentiated colonies is a consequence of the generic proliferative defects of loss of LSD1 as ablation of *Lsd1* did not affect the expression of the pluripotency factors OCT4 and NANOG (**Figure 1I**). Moreover, the immunostaining assay of OCT4 and SSEA did not display any difference in the expression levels between *Lsd1* KO and WT ESCs (**Figure 1J**). Hence, loss of *Lsd1* in ESCs appears to impair its basic proliferative functions while preserving the pluripotent potential.

To identify the transcriptional program of these *Lsd1* KO ESCs, we performed RNA-sequencing (RNA-seq) analysis. Gene expression profiles of both clones were slightly distinct due to normal stochastic heterogeneity between isolated clones (**Figures 1K** and **1L; S1H** and **S1I; Table S1**), yet, KO transcriptomes clustered together and displayed more similarity among them than with the RNA-seq signals retrieved from WT ESCs (**Figure 1M**). Gene ontology (GO) analysis of biological processes of common upregulated genes revealed categories related to generic functions, including negative regulation of cell differentiation, apoptosis, and endodermal differentiation (**Figure 1N**). Common downregulated genes in KO1 and KO2 showed enrichment for nucleosome assembly, DNA repair and metabolic processes, among others (**Figure 1O**). There were no differences in the expression of multiple pluripotency and germ layer-specific markers between WT and *Lsd1* KO ESCs. To validate the RNA-seq data, we performed RT-qPCR on upregulated genes representative of the endoderm such as *Sox17* and *Gata6* (**Figure S1J**). Additionally, we validated the defective metabolic phenotype of *Lsd1* KO ESCs by a flux analyzer. Loss of *Lsd1* led to decreased basal glycolysis and KOs were not able to respond to increased energetic demand even when oxidative phosphorylation was shut down (**Figures 1P** and **1Q**). To assess whether defective glycolysis results in the shift to oxidative metabolism, we performed a mito stress test. However, we did not observe differences in mitochondrial function and ATP production between WT and *Lsd1* KOs, indicating that *Lsd1* KO ESCs might use alternative fuels such as fatty acids to drive OXPHOS-ATP production (**Figures 1R** and **1S**). Taken together, RNA-seq showed that loss of LSD1 does not have a profound effect on the expression of pluripotency-associated genes and thereby, we were able to maintain *Lsd1* KO ESCs in culture over long periods of time without loss of self-renewal.

### Ablation of *Lsd1* leads to defective differentiation

To test the role of LSD1 in differentiation, WT and *Lsd1* KO ESCs were differentiated to EBs for 8 days. *Lsd1* KO showed a significant reduction in the size of EBs compared to EBs derived from WT ESCs (**Figures S2A** and **S2B**), suggesting a defective differentiation phenotype upon loss of *Lsd1*. In addition, *Lsd1* KO ESCs were not able to suppress the expression of the core pluripotency factors *Oct4* and *Nanog* during the course of differentiation (**Figure S2C**). Moreover, EBs generated from both *Lsd1* KO ESCs failed to express lineage-specific markers such as *Sox17* and *Foxa2* (endodermal) (**Figure S2D**), *Brachyury* or *T* and *Msx1* (mesodermal) (**Figure S2E**), and *Sox11* (ectodermal) (**Figure S2F**). In order to exclude potential off-target effects, we next engineered a cell line in which *Lsd1* KO2 ESCs were stably expressing a MYC-tagged WT LSD1 (LSD1^WT^) or a MYC-tagged mutated LSD1 (LSD1^MUT^) (**Figure S2G**). The residues required for demethylating histone H3K4 have not been described for the mouse LSD1 isoform. However, previous studies have shown that double point mutation of human LSD1 at the residues Alanine 539 and Lysine 661 (A539/K661) is required to abolish its demethylase activity [28]. Therefore, we performed multi-species LSD1 protein alignment and we found that A539 and K661 are highly conserved among the represented species (**Figure 2A**). Based on this alignment, we engineered and purified a double point mutated (A540E/K662A) mouse LSD1 protein (LSD1^MUT^) and assayed its activity together with a WT LSD1 (LSD1^WT^). The demethylase activity of LSD1^WT^ was 1.74 ± 0.023 Kcat/ min whereas the enzymatic activity of LSD1^MUT^ was undetectable, indicating that A540E/K662A resulted in a catalytically-dead LSD1 (**Figure 2B**, left and right panel). Strikingly, both LSD1^WT^ and LSD1^MUT^ cell lines rescued the proliferation defect of *Lsd1* KO ESCs and showed colony morphology and AP staining comparable to WT ESCs (**Figures S2H-J**). However, the defective size of *Lsd1* KO EBs was only partially recovered in LSD1^MUT^ (**Figures 2C** and **2D**), as LSD1^MUT^ EBs were not able to silence the core pluripotency factors and to enhance the expression of lineage markers similar to *Lsd1* KO EBs (**Figures 2E-2I**). Overall, these results indicate that albeit the enzymatic activity of LSD1 is not required to maintain the proliferation capacity of mouse ESCs, it is essential to trigger proper differentiation.

**Figure 2.**
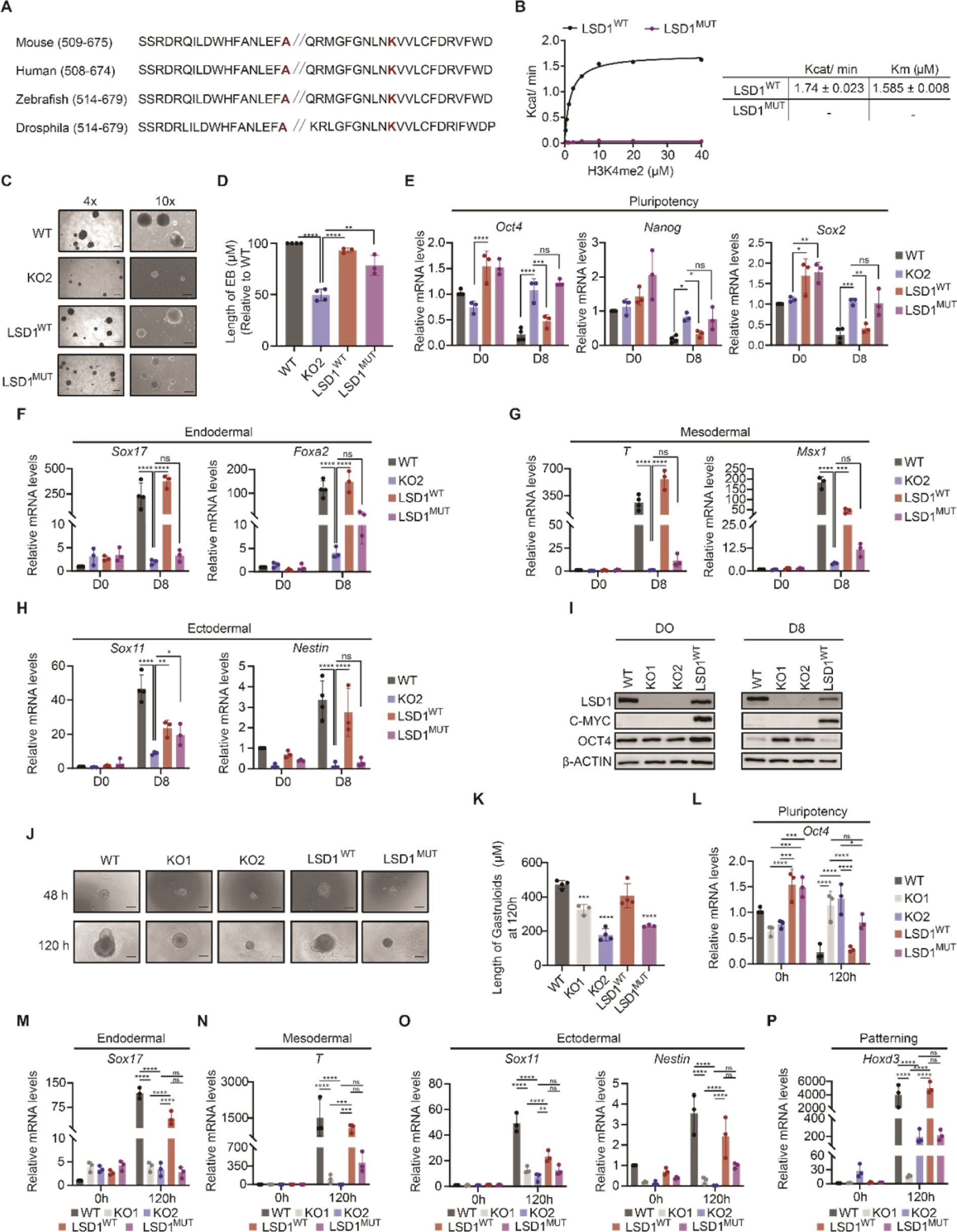
**LSD1 is essential for ESC differentiation.** (A) Sequence alignment of LSD1 in different species. The amino acid residues that were pointed mutated are highlighted in red. (B) Saturation curves of histone demethylase activity of purified WT and mutant LSD1 proteins with increasing concentrations of H3K4me2 as a substrate. Each enzymatic curve was derived from the Michaelis-Menten equation (left panel) and their reaction kinetics (right panel). (C and D) (C) Representative bright field images at (4x (left) and 10x (right)) magnification (D) quantification of the size of EB derived from WT, *Lsd1* KO2, LSD1^WT^, and LSD1^MUT^ ESCs at day 8 of differentiation. Scale bars, 200 µM. (E-H) RT-qPCR analysis of (E) pluripotency (*Oct4*, *Nanog,* and *Sox2*), (F) endodermal (*Sox17* and *Foxa2*), (G) mesodermal (*T* and *Msx1*), and (H) ectodermal (*Sox11* and *Nestin*) markers in WT, *Lsd1* KO2, LSD1^WT,^ and LSD1^MUT^ ESCs on day 0 and day 8 of EB differentiation. mRNA levels are relative to the expression of WT at day 0. *β-actin* is used as an internal control. (I) Western blot of LSD1, C-MYC, and OCT4 on whole cell extract of EBs of WT, *Lsd1* KO1, *Lsd1* KO2, and LSD1^WT^ ESCs at day 0 and day 8 of EB differentiation. β-ACTIN is used as the loading control. (J and K) (J) Morphological representation (10x magnification) and (K) measurement of the length of gastruloids derived from WT, *Lsd1* KO1, *Lsd1* KO2, LSD1^WT,^ and LSD1^MUT^ ESCs at indicated time points. Scale bar, 200 µM. (L-P) Bar graph depicting the RT-qPCR analysis of (L) pluripotency (*Oct4*), (M) endodermal (*Sox17*), (N) mesodermal (*T*), (O) ectodermal (*Sox11* and *Nestin*), and (P) patterning (*Hoxd3*) markers in the gastruloids generated from WT, *Lsd1* KO1, *Lsd1* KO2, LSD1^WT,^ and LSD1^MUT^ ESCs at 0h and 120h of gastrulation. mRNA levels are relative to the expression of WT at 0h. *β-actin* is used as an internal control. Statistical analysis: unpaired t-test (D, J) and two-way ANOVA (E-H and L-P). ∗p < 0.05, ∗∗p < 0.01, ∗∗∗p < 0.001, and ∗∗∗∗p < 0.0001. Error bars denote mean ± SD; n ≥ 3 (D-H and K-P). Results are one representative of n ≥ 3 independent experiments (C, I and J).

To validate our findings that loss of LSD1 impairs ESC differentiation, we tested the role of LSD1 in gastruloid generation. Gastruloids are small aggregates of ESCs that, under appropriate culture conditions, recapitulate the axial organisation of post-implantation embryos and mimic major aspects of gastrulation [29]. *Lsd1*/LSD1 was uniformly expressed, implying that LSD1 has functional importance during gastrulation, whereas the expression of *Oct4*/OCT4 was decreased after 120h of gastruloid formation (**Figures S2K** and **S2L**). We next generated gastruloids from WT, *Lsd1* KO ESCs, LSD1^WT^ and LSD1^MUT^ ESCs. *Lsd1* KO ESCs failed to give rise to elongated gastruloid-like structures, being these aggregates significantly smaller than in WT ESCs (**Figures 2J** and **2K**). Noteworthy, the morphologic phenotype was rescued in LSD1^WT^ but not in LSD1^MUT^ ESCs (**Figures 2J** and **2K**). Similar to what we observed during EB formation, gastruloids derived from *Lsd1* KO and LSD1^MUT^ ESCs retained higher levels of *Oct4* compared with WT, and the expression of the differentiation markers and the axial patterning marker *Hoxd3* failed to be upregulated (**Figures 2L-P**). Collectively, these results show that loss of LSD1 abrogates proper gastruloid formation and revealed that the catalytic activity of LSD1 is required for proper ESC differentiation.

### Loss of LSD1 results in accumulation of H3K4me1

To gain a better overview of the LSD1-mediated regulation of the histone modification landscape in mouse ESCs, we performed nanoscale liquid chromatography coupled to tandem mass spectrometry (nano LC-MS/MS), a robust quantitative method for the characterization of post-translational modifications (PTMs) of histones. Because a given PTM is commonly present in different peptides due to combinatorial PTM possibilities, we focused on the analysis of deconvoluted single PTMs. Hence, to retrieve the relative abundance of histone PTMs, the sum of all modified forms in their respective histone peptides was considered as 100% (**Table S2**). After calculating the co-existence of the individual histone marks in WT ESCs, our analysis revealed that H3K27me2 and H3K36me2 were the two most abundant methylation marks as previously reported, whereas most of K4, K14, K18 and K23 were not modified on histone H3 (**Figure 3A**) [30]. Next, we compared the relative abundance of histone marks in WT and *Lsd1* KO ESCs (**Figure S3A**). H3_3_8K4me1 was the most abundant peptide in both KOs compared to WT ESCs (**Figures S3B** and **S3C**). Moreover, our analysis showed that H3K4me1 was significantly increased upon loss of LSD1, whereas no differences were observed in H3K4me2 and H3K4me3 between WT and *Lsd1* KO ESCs (**Figure 3B**). Strikingly, no changes in H3K9me1 were observed between WT and *Lsd1* KO ESCs, whilst the latest displayed a decrease of H3K9me2 and H3K9me3 marks compared to WT (**Figure 3C**). This data indicates that, in mouse ESCs, LSD1 demethylates the active histone modification mark H3K4me1 thereby acting as a transcriptional repressor.

**Figure 3.**
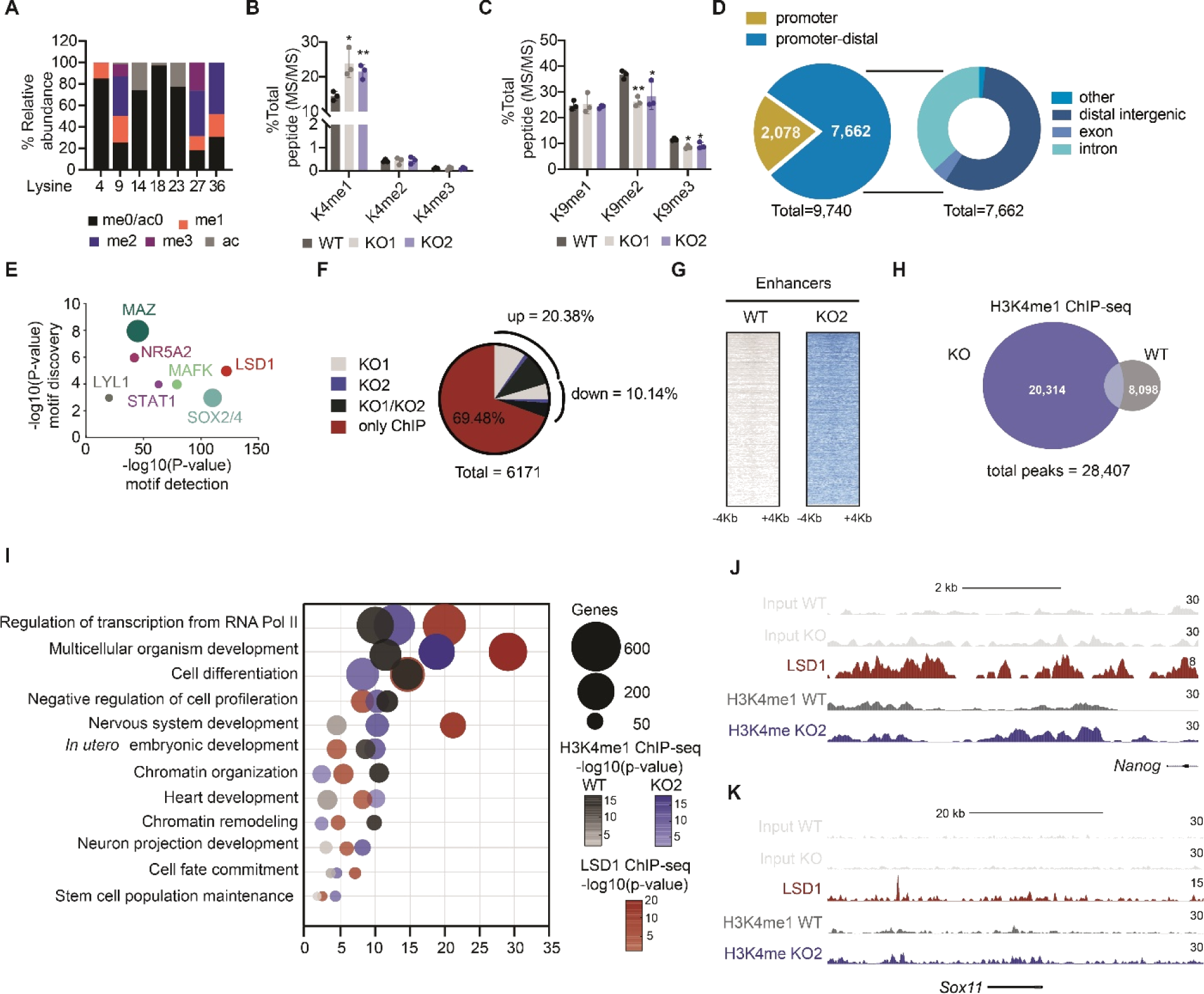
**Ablation of LSD1 affects global H3K4me1 levels.** (A) Relative abundance of histone marks in different positions of lysine on the N-terminal tail of histone H3 in WT mouse ESCs. The colour coding represents different post-translational modifications (PTM), and the y-axis represents their relative abundance. (B and C) (B) Quantification of H3K4 methylation and (C) H3K9 methylation levels in WT and *Lsd1* KO ESCs. Each modification represents the percentage of one modified peptide among the total modified peptides observed in independent replicates. (D) Genomic distribution of LSD1 binding in the promoter (within 5 kb upstream of TSS) and promoter-distal regions (left) and the fraction of different regions such as distal intergenic, exon, and intron within promoter-distal regions (right) in WT mouse ESCs. (E) Enriched transcription factor motifs at LSD1 peaks in WT mouse ESCs. (F) Pie chart representing the percentage of LSD1-only bound genes, bound and activated in *Lsd1* KO1, KO2 or KO1/KO2 ESCs, and bound and repressed in *Lsd1* KO1, KO2 or KO1/KO2 ESCs. (G) Density plots of H3K4me1 ChIP-seq at the enhancer regions in WT and *Lsd1* KO2 ESCs. (H) Venn diagram of overlapped genes in H3K4me1 ChIP-seq between WT and *Lsd1* KO2 ESCs. (I) GO analysis of biological processes of genes associated with LSD1 peaks in WT ESCs and H3K4me1 peaks in WT and *Lsd1* KO2 ESCs. (J and K) LSD1 ChIP-seq signal in mouse ESCs and H3K4me1 ChIP-seq signal in WT and *Lsd1* KO2 ESCs at the (J) *Nanog* and (K) *Sox11* enhancers. Respective inputs are depicted in grey. Statistical analysis: unpaired t-test (B-C). ∗p < 0.05, ∗∗p < 0.01. Error bars denote mean ± SD; n=3 (B-C).

To further investigate H3K4me1 levels regulated by LSD1, we performed chromatin immunoprecipitation-coupled with high-throughput DNA massively parallel sequencing (ChIP-seq) experiments (ChIP-seq) with antibodies against either H3K4me1 or LSD1. We identified a total of 9,740 LSD1 peaks throughout the genome (**Figure 3D; Table S3**). Of these, the vast majority were promoter-distal (7,662; >3 kb from TSS), which is consistent with prior observations [16]. LSD1 peaks overlapped with binding sites for transcription factors with a reported stem cell function such as MAZ, NR5A2, and Sox2/4, and other transcriptional factors involved in the differentiation of ESCs (**Figure 3E**) [16, 31, 32]. Integration of RNA-seq in WT and *Lsd1* KO ESCs with ChIP-seq data showed that the majority of differentially expressed genes are not direct targets of LSD1 (∼70%; **Figure 3F**). We identified a total of 28,407 H3K4me1 peaks which mostly decorated enhancer regions (**Figure 3G; Table S4)**. In addition, ∼71 % and ∼28 % of those sites gained H3K4me1 in *Lsd1* KO and WT ESCs, respectively, consistent with the observation that loss of LSD1 leads to an accumulation of H3K4me1 in mouse ESCs (**Figure 3H**). We found that global increase of H3K4me1 in *Lsd1* KO ESCs did not correlate with major changes in gene expression (**Figure S3D**). Particularly, 1414 genomic regions gaining H3K4me1 showed a significant upregulation of the expression of transcripts whereas only a minority (328) displayed downregulation (**Figure S3D**). GO analysis for biological processes showed that LSD1 binds preferentially to genes related to ‘regulation of transcription’, ‘multicellular organism development’ and ‘cell differentiation’ (**Figure 3I**). Likewise, H3K4me1 peaks that were gained in both WT and *Lsd1* KO ESCs were enriched in similar categories (**Figure 3I**). We also found binding of LSD1 at genes important for ‘Stem cell population maintenance*’* although to a lesser extent (**Figure 3I**). We next identified genes that are repressed (1588) or overexpressed (462) during RA-mediated differentiation (FC > 1.5) by using publicly available data [33]. When overlapping the aforementioned genes with LSD1 ChIP-seq, we found that both induced and repressed genes were bound by LSD1 (**Figure S3E**), suggesting that LSD1 does not promote or inhibit stem cell fate but instructs a more general role in gene expression of mouse ESCs [34]. Hence, enhancers controlling the expression of both pluripotency (e.g. *Nanog*, *Prdm14*, *Oct4*) and developmental genes (e.g. *Sox11* and *Sox17*) gained H3K4me1 upon loss of LSD1 (**Figures 3J-K** and **S3F-H**).

### Loss of LSD1 leads to global DNA hypomethylation

It has been shown that targeted deletion of LSD1 leads to progressive loss of global DNA methylation [6, 20, 23]. Hence, to further explore the role of LSD1 on DNA methylation, we employed liquid chromatography-tandem mass spectrometry (LC-MS/MS) and measured the abundance of 5-methylcytosine (5mC) and the oxidized 5mC derivative, 5-hydroxymethylcytosine (5hmC), in genomic DNA of WT and *Lsd1* KO mouse ESCs. Our analysis revealed that both 5mC and 5hmC levels were remarkably decreased upon loss of *Lsd1* (**Figures 4A** and **S4A**). Dot blot on isolated genomic DNA and immunofluorescence analysis with specific antibodies against 5mC and 5hmC confirmed the same result (**Figures 4B, S4B** and **S4C**). The 5hmC/5mC ratio remained similar between WT and *Lsd1* KO ESCs, suggesting that lower levels of 5hmC are a consequence of the reduced amount of available 5mC substrate rather than active 5mC hydroxylation (**Figure S4D**). Next, in order to exclude potential off-target effects, we assessed global DNA methylation levels in LSD1^WT^ ESCs. LSD1^WT^ ESCs exhibited 5mC levels similar to WT ESCs whilst re-introduction of an empty MYC vector (KO2^EV^) was not able to rescue the hypomethylation phenotype observed upon *Lsd1* loss (**Figures 4C** and **4D**). DNA hypomethylation levels of *Lsd1* KO ESCs were similar to *Dnmt1* KO ESCs, but relatively higher than *Dnmt3a/Dnmt3b* double KO ESC lines (**Figures S4E** and **S4F**). Remarkably, LSD1^MUT^ ESCs exhibited comparable DNA methylation levels to WT and LSD1^WT^ ESCs (**Figure 4E**), indicating that, in contrast to previous studies [6, 20], the catalytic activity of LSD1 does not play a role in DNA methylation.

**Figure 4.**
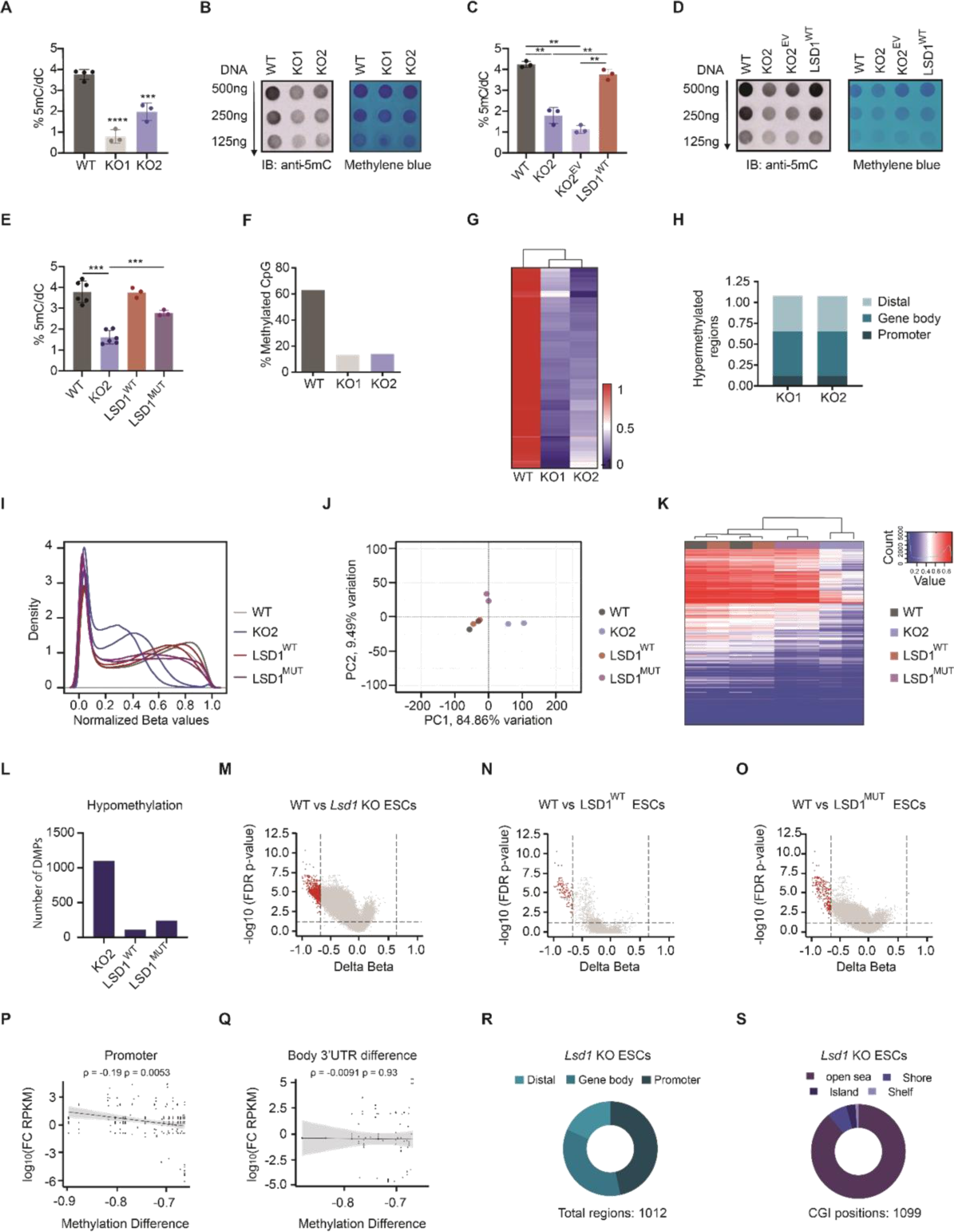
**Loss of LSD1 affects global DNA methylation.** (A and B) (A) LC-MS/MS quantification of 5mC/dC and (B) dot blot analysis of 5mC/dC (left panel) of genomic DNA extracted from WT and *Lsd1* KO ESCs. Methylene blue staining was used as loading control (right panel). (C and D) (C) LC/MS-MS quantification and (D) dot blot analysis of 5mC/dC (left panel) of genomic DNA extracted from WT, *Lsd1* KO2, KO2^EV^, and LSD1^WT^ ESCs. Methylene blue staining was used as loading control (right panel). (E) LC/MS-MS quantification of genomic DNA extracted from WT, *Lsd1* KO2, LSD1^WT,^ and LSD1^MUT^ ESCs. (F) Bar graph showing percentage methylated CpGs in *Lsd1* KO ESCs and WT. (G) Heatmap generated from whole-genome bisulfite-sequencing depicting global DNA methylation in WT and *Lsd1* KO ESCs. Red corresponds to the hypermethylation CpG sites, and in blue, the hypomethylation CpG sites. (H) Percentage of hypomethylation regions in the promoter, gene body, and distal regulatory elements in *Lsd1* KOs. Promoter (-5kb to +500bp from TSS), Gene body (+500 bp from TSS to +500bp from TES), and Distal (>5kb upstream or >500bp downstream) were considered for the analysis. (I) Density plot of DNA methylation β values of WT, *Lsd1* KO2, LSD1^WT,^ and LSD1^MUT^ ESCs. Methylation values range from zero (fully unmethylated) to one (fully methylated). (J) Principal component analysis of array-based DNA methylation profiles of WT, *Lsd1* KO2, LSD1^WT,^ and LSD1^MUT^ ESCs. (K) Heatmap and unsupervised hierarchical clustering of methylation levels in 10,000 random CpGs in WT, *Lsd1* KO2, LSD1^WT,^ and LSD1^MUT^ ESCs. Red corresponds to the hypermethylated CpG sites, and blue to the hypomethylated CpG sites. Hierarchical clustering was performed with Euclidean distance and Ward’s minimum variance agglomeration method. (L) Bar graph representing number of differentially hypomethylated positions in *Lsd1* KO2, LSD1^WT^, and LSD1^MUT^ compared to WT ESCs. (M-O) Volcano plots of differentially methylated positions (DMPs) in (M) *Lsd1* KO2 (N) LSD1^WT^ and (O) LSD1^MUT^ compared to WT ESCs. Red dots indicate significant results (FDR < 0.05 and value of Δβ < -0.66 or >0.66). (P and Q) Scatter correlation plots of (P) promoter methylation and (Q) body or 3’UTR methylation and gene expression in *Lsd1* KO compared to WT ESCs. Y-axis represents the log10 fold change of Reads Per Kilobase of transcript per Million mapped reads (FC RPKM) from RNA-seq and x-axis represents β -value methylation difference. Spearman rank correlation coefficient and corresponding p-value are shown. (R) Genomic distribution of hypomethylated regions in *Lsd1* KO2 compared to WT ESCs. (S) Pie chart representing distribution of hypomethylated sites with respect to CGI positions. In *Lsd1* KO2 compared to WT ESCs Statistical analysis: unpaired t-test (A, C, and E). ∗∗p < 0.01, ∗∗∗p < 0.001, and ∗∗∗∗p < 0.0001. Error bars denote mean ± SD; n ≥ 3 (A, C and E). Results are one representative of n = 3 independent experiments (B and D).

To further evaluate the role of LSD1 in DNA methylation, we next assessed the DNA methylation landscape by whole-genome bisulfite sequencing. The genome was divided into 1 kb tiles, and the distribution of CpG methylation values was compared within each tile across the samples. We identified 83,671 and 83,950 differentially methylated regions (DMRs) in *Lsd1* KO1 and KO2 compared to WT, respectively, confirming the hypomethylated phenotype upon loss of LSD1 (**Figures 4F** and **4G**). To gain a global perspective on the effect of DNA methylation on CpG islands and their neighbouring regions, we generated composite plots for mean methylation levels across 25,489 CpG islands. We found a general decrease in methylation in CpG islands which was not just at the centre of the island but also on CpG island shores (**Figure S4G**). Most of the DMR were enriched for gene bodies (+500 bp from the transcription start site (TSS) to +500 bp from the transcription end site (TES)) and distal regions (> 5kb upstream or > 500 bp downstream) whereas hypomethylated promoters (-5kb to +500bp from TSS) were underrepresented (**Figure 4H**). Such DMR distal regions were enriched for repetitive elements such as SINEs and LINEs, and for enhancer regions, whereas we found a quantitative association between DNA methylation at promoters of WT and *Lsd1* KO ESCs, confirming our previous observation that promoter sequences do not suffer major changes in methylation upon loss of LSD1 (**Figures S4H-S4J**). Altogether, these data suggest that LSD1 regulates DNA methylation genome-wide, especially in repetitive elements and enhancers. We next assessed the methylation levels of known imprinted genes as genome imprinting is regulated by 5mC deposition. Particularly, the paternally imprinted H19 and insulin-like growth factor (Igf2) gene cluster, the intergenic differentially methylated region (IG-DMR) which regulates the allele-specific expression of the *Dlk1-Dio3* cluster, the *Kcnq1ot1* imprinting control region and the maternal imprinted gene *Peg3* underwent hypomethylation in *Lsd1* KO ESCs (**Figure S4K**). Hence, our data indicates that LSD1 plays an important role in mediating methylation of imprinted genes.

To further understand the function of LSD1 in DNA methylation, we profiled 5mC levels in WT, *Lsd1* KO, LSD1^WT^ and LSD1^MUT^ ESCs by employing the Mouse Methylation MM285 BeadChIP microarray [35]. We defined differentially methylated CpG sites as those with absolute methylation difference >66% and adjusted p-value < 0.05 (**Table S5**). DNA methylation β-value density plots showed a bimodal distribution, being the unmethylated peak higher than the methylated peak, reflecting a genome-wide DNA methylation loss that mouse ESCs exhibit (**Figure 4I**). Furthermore, the number of fully methylated CpGs was lower for *Lsd1* KO ESCs compared to the other cell lines analyzed. Indeed, principal component analysis showed a clear separation between the WT and *Lsd1* KO ESCs, whereas both LSD1^WT^ and LSD1^MUT^ clustered close to WT ESCs, confirming that the hypomethylation phenotype of *Lsd1* KO ESCs is rescued upon the introduction of the catalytically dead LSD1 (**Figure 4J**). Unsupervised analysis of 10,000 random CpGs revealed the same finding with similar methylation patterns in WT, LSD1^WT^, and LSD1^MUT^, but distinct from the *Lsd1* KO ESCs (**Figure 4K**). We identified a set of 1,099 loci that were hypomethylated upon LSD1 loss, whereas only 111 and 240 were hypomethylated in LSD1^WT^ and LSD1^MUT^ ESCs, respectively, compared to WT ESCs (**Figures 4L-O**). Specifically, we found 111 probes in LSD1^WT^ ESCs, corresponding to 10 genes, that remained hypomethylated upon reintroduction of a WT LSD1, suggesting intrinsic differences amongst cell lines. Those genes remained hypomethylated also in LSD1^MUT^ ESCs. In addition, 126 loci, corresponding to 46 genes, were hypomethylated in *Lsd1* KO and LSD1^MUT^ ESCs, indicating that the catalytic activity of LSD1 is required to establish proper methylation at some specific loci (**Figures S4L** and **S4M**). Of note, we only found a negligible number of hypermethylated genes in LSD1^MUT^ ESCs. Differential methylation between WT and *Lsd1* KO ESCs at promoter regions, but not at body or 3’UTR, negatively correlated with changes in gene expression (**Figures 4P** and **4Q; Table S6**). We next explored the genomic distribution of hypomethylated genomic elements from *Lsd1* KO ESCs. Intriguingly, promoter regions (corresponding to TSS 1500, TSS 200, 5′UTR and 1st Exon) were more represented than in the bisulfite analysis, albeit we also detected hypomethylation in the gene bodies and intergenic regions (**Figure 4R**). Most of the CpGs identified were significantly enriched in open sea regions (89.08 %) and only a minority of them associated with (3.18%) CpG islands, (1.00%) with shelves (2-4 kb from the promoter CpG islands) and (6.73%) with shores (0-2 kb from the promoter CpG islands) (**Figure 4S**). Altogether, this data show that loss of LSD1 leads to global hypomethylation, and that this effect is mostly independent of LSD1 demethylase activity.

### LSD1 catalytic activity is not required for DNMT1 and UHRF1 expression

Because several studies have shown that LSD1 stabilises both DNMT1 and UHRF1 proteins [6, 33], we next assessed whether the protein expression levels of the DNA maintenance methylation machinery were altered in *Lsd1* KO ESCs. Western blot analysis on whole cell and chromatin extracts showed reduced DNMT1 and UHRF1 upon *Lsd1* loss **(Figures 5A** and **5B)**. Such decreased of DNMT1 and UHRF1 proteins were not associated with changes in mRNA levels as indicated by our RNA-seq analysis (**Figures S5A** and **S5B**). However, the expression of *de novo* DNA methyltransferases DNMT3A and DNMT3B, and the accessory factor DNMT3L, were increased in *Lsd1* KO compared to WT ESCs, albeit exhibiting lower global 5mC levels than WT (**Figures S5C** and **S5D**). In this case, protein abundances correlated with mRNA expression levels (**Figures S5E-S5G**). Similarly, DNMT3A, DNMT3B, and DNMT3L proteins were also significantly increased in *Dnmt1* KO ESCs (**Figures S5H** and **S5I**), suggesting that activated expression of *de novo* methyltransferases is a compensatory mechanism to counteract the loss of DNMT1 [36]. Reintroduction of LSD1 in *Lsd1* KO ESCs rescued the DNMT1 and UHRF1 protein levels (**Figures 5C** and **5D**). In agreement with the observation that concomitant expression of mutated LSD1 in *Lsd1* KO ESC recovered the hypomethylation phenotype (**Figure 4E**), LSD1^MUT^ ESCs were also able to restore DNMT1 and UHRF1 protein abundance without affecting *Dnmt1* and *Uhrf1* mRNA expression levels **(Figures 5C-5F).** Altogether, and in contrast to previous studies, our data suggest that LSD1 sustains DNMT1 and UHRF1 protein stability independent of the demethylase activity.

**Figure 5.**
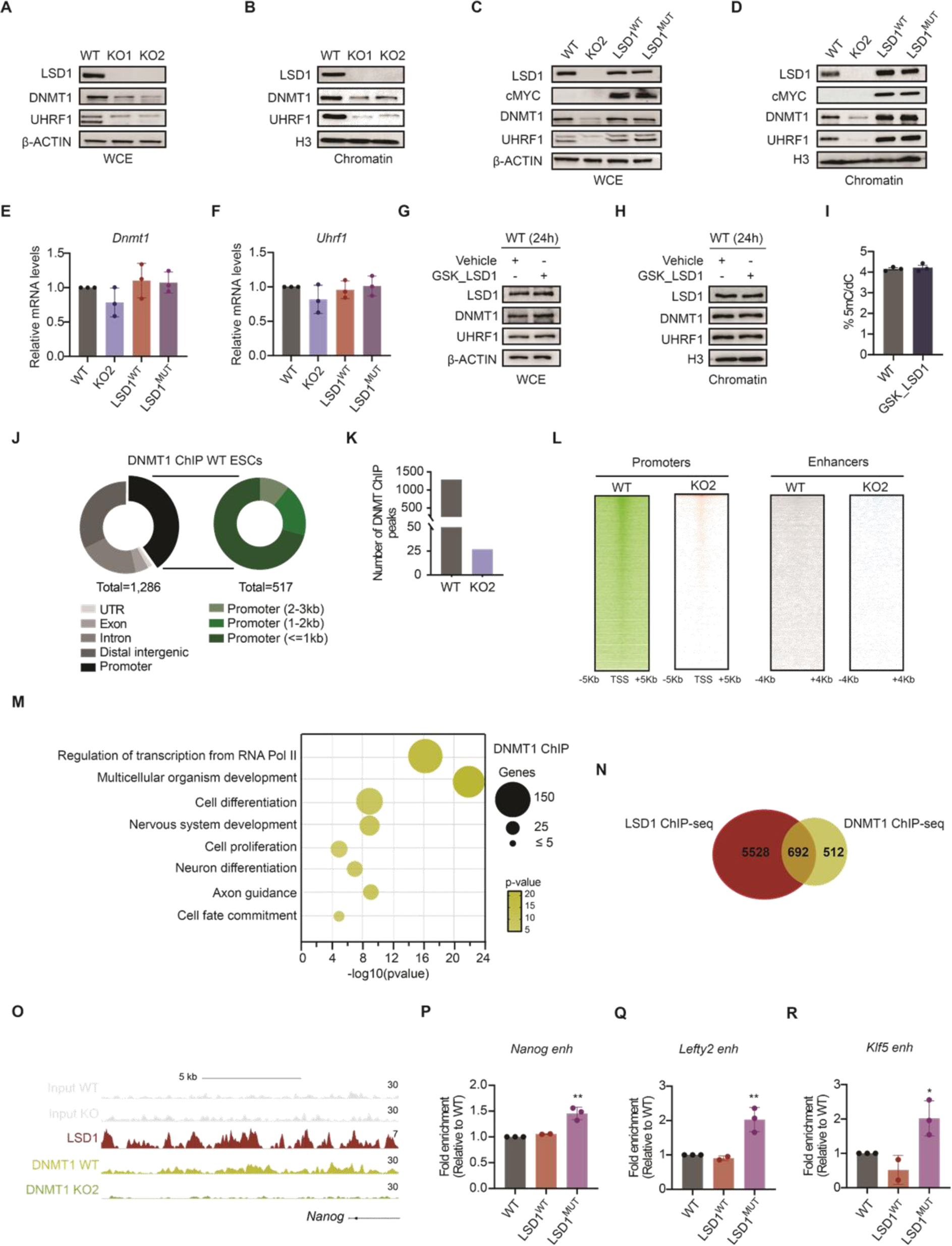
**Deletion of *Lsd1* leads to impaired DNA methylation machinery.** (A and B) Western blots of LSD1, DNMT1, and UHRF1 on the (A) WCE and (B) chromatin fractions of WT and *Lsd1* KO ESCs. β-ACTIN and H3 are used as the loading controls. (C and D) Western blotting assay with antibodies against LSD1, cMYC, DNMT1, and UHRF1 on (C) WCE and (D) chromatin fractions of WT, *Lsd1* KO2, LSD1^WT,^ and LSD1^MUT^ ESCs. β-ACTIN and H3 are used as the loading controls. (E and F) Relative levels of *Dnmt1* and *Uhrf1* mRNA in the WT, *Lsd1* KO2, LSD1^WT^, and LSD1^MUT^ ESCs assessed by RT-qPCR. (G and H) Western blots of LSD1, DNMT1 and UHRF1 on (G) WCE and (H) chromatin fractions of vehicle and LSD1 inhibitor (GSK_LSD1) treated WT ESCs. β-ACTIN and H3 are used as the loading controls. (I) LC-MS/MS quantification of 5mC on genomic DNA in WT mouse ESCs treated with a vehicle and inhibitor (GSK_LSD1). (J) Genomic distribution of DNMT1 binding regions (left) and within different regions of the promoter (right). (K) Bar diagram depicting number of DNMT1 ChIP peaks in WT and *Lsd1* KO2 ESCs. (L) DNMT1 ChIP–seq heatmap in WT and *Lsd1* KO2 ESCs. (M) GO analysis of biological processes of genes associated with DNMT1 peaks in WT mouse ESCs. (N) Venn diagram of overlapped genes associated with LSD1 and DNMT1 ChIP-seq peaks in WT ESCs. (O) LSD1 ChIP-seq signal in mouse ESCs and DNMT1 ChIP-seq signal in WT and *Lsd1* KO2 ESCs at the *Nanog* enhancer. Respective inputs are depicted in grey. (P-R) ReChIP-qPCR analysis of DNMT1 enrichment at (P) *Nanog,* (Q) *Lefty2,* and (R) *Klf5* loci in WT, LSD1^WT,^ and LSD1^MUT^ ESCs. The data were normalized to input and represented relative to respective WT. Statistical analysis: ordinary one-way ANOVA (E and F) and unpaired t-test (I and P-R). ∗p < 0.05, ∗∗p < 0.01. Error bars denote mean ± SD; n ≥ 2 (E-F, I and P-R). Results are one representative of n = 3 independent experiments (A-D and G-H).

To further investigate whether the catalytic activity of LSD1 is dispensable for the expression of DNMT1 and UHRF1, we treated ESCs with GSK-LSD1, a chemical LSD1 inhibitor which has been shown to be involved in the covalent modification of the cofactor flavin adenine dinucleotide (FAD) [37]. The efficiency of the LSD1 inhibitor was determined by performing RT-qPCR on direct targets of LSD1, such as *Sox17* and *Eomes* **(Figures S5J** and **S5K)**. Noteworthy, GSK-LSD1 treatment did not affect *Lsd1*, *Dnmt1* and *Uhrf1* mRNAs nor LSD1, DNMT1 or UHRF1 protein levels (**Figures 5G, 5H** and **S5L-S5N)**. Furthermore, GSK_LSD1 treated ESCs exhibited 5mC levels similar to WT ESCs (**Figure 5I)**, confirming that the catalytic activity of LSD1 is not required for DNMT1 and UHRF1 expression, and DNA methylation maintenance.

Given that DNMT1 protein levels were drastically reduced upon loss of LSD1, we aimed to map DNMT1 binding by ChIP-seq in WT and in *Lsd1* KO ESCs. We found 1,286 DNMT1 peaks that were distributed along the genome (**Figure 5J; Table S7**). Consistent with decreased DNMT1 protein abundance upon LSD1 loss, we only recovered 27 peaks in *Lsd1* KO ESCs (**Figure 5K**). The density of DNMT1 signals was higher at the promoters than enhancers in WT ESCs, whereas no distinct signal was observed in *Lsd1* KO ESCs (**Figure 5L).** GO analysis for biological processes indicated that DNMT1 binds preferentially to genes enriched in multicellular organism development, regulation of transcription, and neuron-related categories (**Figure 5M**). Around 55 % of these sites were co-occupied by LSD1, suggesting the possibility that, besides regulating DNMT1 stability, LSD1 could be also involved in DNMT1 chromatin recruitment (**Figure 5N**). Hence, to further confirm LSD1 dependency of DNMT1 chromatin binding we performed re-ChIP followed by RT-qPCR of common LSD1 and DNMT1 overlapping regions **(Figures 5O, S5O** and **S5P**). Thus, following immunoprecipitation with the first ChIP LSD1 antibody, LSD1-bound fragments underwent the second ChIP with antibodies against DNMT1 in WT, LSD1^WT^ and LSD1^MUT^ ESCs. We found DNMT1 present in *Nanog*, *Lefty2* and *Klf5* enhancers containing both WT and LSD1^MUT^, indicating that LSD1 acts as a scaffold protein to recruit DNMT1 at specific loci (**Figures 5P-R**).

### The scaffolding function of LSD1 stabilises DNMT1 and UHRF1 proteins

The observation that LSD1 regulated DNMT1 and UHRF1 protein levels indicated that LSD1 controls DNMT1 and UHRF1 stability. Thus, we first analysed whether LSD1 interacts with DNMT1 and UHRF1. To this aim, we performed co-immunoprecipitation experiments in whole cell extracts and nuclear fractions of WT, *Lsd1* KO, and LSD1^MUT^ ESCs. We detected interaction of DNMT1 and UHRF1 with LSD1 in both WT and LSD1^MUT^ ESCs (**Figures 6A** and **6B)**. We next examined whether DNMT1 and UHRF1 protein stability was affected in the absence of LSD1 or when the catalytic centre of LSD1 was mutated. To this end, we treated WT, *Lsd1* KO, LSD1^WT^, and LSD1^MUT^ ESCs with protein synthesis inhibitor cycloheximide (CHX) and then monitored the rates of DNMT1 and UHRF1 decline over time. Our data revealed that DNMT1 and UHRF1 half-life was dramatically decreased upon loss of LSD1. However, DNMT1 and UHRF1 had comparable rates of degradation and half-lives in WT, LSD1^WT^, and LSD1^MUT^ ESCs, implying that the demethylase activity of LSD1 is not required to sustain DNMT1 and UHRF1 protein stability **(Figures S6A** and **6C-6D**).

**Figure 6.**
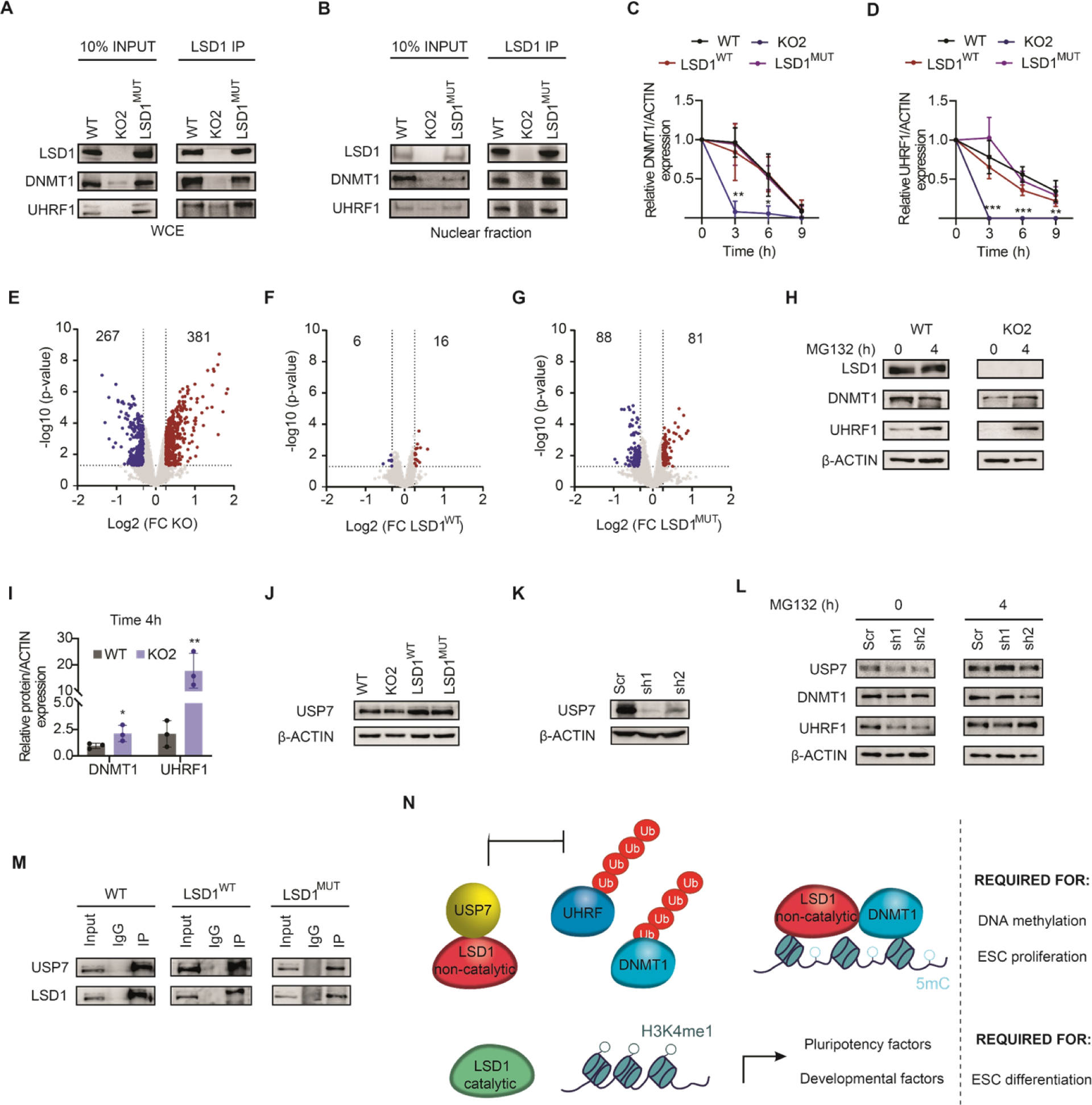
**Loss of LSD1 diminishes DNMT1 and UHRF1 stability.** (A and B) LSD1 immunoprecipitation in (A) WCE and (B) nuclear fraction of WT, *Lsd1* KO2, and LSD1^MUT^ ESCs followed by immunoblotting of LSD1, DNMT1 and UHRF1. The percentage of input used is 10%. (C and D) Protein degradation curves of (C) DNMT1 and (D) UHRF1 were generated from the quantification and normalization of the bands from (S6A) upon CHX treatment. (E-G) Volcano plots of differentially expressed proteins in (E) *Lsd1* KO2, (F) LSD1^WT^, and (G) LSD1^MUT^ ESCs in comparison to WT ESCs. Significant upregulated and downregulated proteins are represented in red and blue, respectively (p < 0.05 and FC (downregulated < 0.8 and upregulated > 1.2)). (H and I) (H) Western blots of LSD1, DNMT1, and UHRF1 in whole cell extract of WT and *Lsd1* KO2, ESCs at 0 and 4 hours after MG132 treatment. (I) Bar graph representing relative protein recovery after quantification and normalization of the bands from (E). β-ACTIN is used as the loading control. (J) Western blot of USP7 in whole cell extract of WT, *Lsd1* KO2, LSD1^WT,^ and LSD1^MUT^ ESCs. β-ACTIN is used as the loading control. (K) Western blot of USP7 in WT ESCs upon scramble and knockdown of *Usp7*. β-ACTIN is used as the loading control. (L) Western blots of USP7, DNMT1, and UHRF1 in WT ESCs upon scramble and knockdown of *Usp7* at 0 and 4 hours after MG132 treatment. β-ACTIN is used as the loading control. (M) USP7 immunoprecipitation in WT, LSD1^WT^, and LSD1^MUT^ ESCs followed by immunoblotting of USP7 and LSD1. The percentage of input used is 10%. (N) Graphical illustration of the model. Non-catalytic LSD1 interacts with USP7 to facilitate USP7-mediated ubiquitination of DNMT1 and UHRF1, promoting DNMT1 and UHRF1 protein stability. Likewise, non-catalytic LSD1 can also recruit DNMT1 at specific loci. Non-catalytic LSD1 is required for DNA methylation and ESC proliferation. LSD1 catalytic activity is required for H3K4me1 demethylation at both pluripotency and developmental enhancers. Such demethylase activity is required for proper ESC differentiation. Statistical analysis: unpaired t-test (C, D and I). ∗p < 0.05, ∗∗p < 0.01, and ∗∗∗p < 0.001. Error bars denote mean ± SD; n = 3 (C, D and F). Results are one representative of n = 3 independent experiments (A, B, H, J, K, L and M).

To further investigate whether LSD1 could mediate global protein expression levels independently of its catalytic activity, we performed proteomics analysis in WT, *Lsd1* KO, LSD1^WT^, and LSD1^MUT^ ESCs using LC-MS/MS (**Table S8**). In *Lsd1* KO ESCs compared to WT ESCs, 267 proteins were downregulated, while 381 proteins were upregulated (**Figure 6E**). These findings indicate that the loss of LSD1 can exert both stabilizing and destabilizing effects on global protein expression as previously reported [25, 26, 38]. Expectedly, LSD1^WT^ ESCs were virtually able to rescue the number of dysregulated proteins (**Figure 6F**). Furthermore, we found that only 88 proteins were downregulated and 81 proteins were upregulated in LSD1^MUT^ ESCs, indicating that the majority of the changes in protein levels observed in *LSD1* KO ESCs are not dependent on LSD1 enzymatic activity (**Figures 6G**). GO analysis of biological processes of downregulated proteins exclusive to KO revealed categories related to translation, chromatin organization, and protein stabilization (**Figures S6B** and **S6C**). Conversely, the upregulated proteins exclusive to the KO cells were enriched in categories such as protein transport and apoptosis (**Figures S6D** and **S6E**).

Given that LSD1 promotes DNMT1 and UHRF1 stability, we next aimed to address whether the ubiquitin-proteasome pathway is involved in mediating DNMT1 and UHRF1 turn over. To this end, we treated WT and *Lsd1* KO ESCs with the proteasome inhibitor MG132. The addition of MG132 significantly recovered UHRF1 and DNMT1 protein levels in *Lsd1* KO ESCs, albeit the effect was more robust in UHRF1 than in DNMT1 (**Figures 6H** and **6I**), suggesting that LSD1 protects DNMT1 and UHRF1 from proteasome-mediated degradation. Previous studies have shown that the deubiquitinating enzyme USP7 plays a role in promoting protein stability of both DNMT1 and UHRF1 [39, 40]. Additionally, a direct interaction of LSD1 with USP7, in which LSD1 is a target of USP7, has also been described [41]. In order to investigate the role of USP7 in our system, we first examined USP7 expression in whole cell extracts of WT, *Lsd1* KO, LSD1^WT^, and LSD1^MUT^ ESCs. We did not find any significant difference in USP7 protein levels amongst the four cell lines analysed (**Figure 6J**). We next sought to determine whether silencing of USP7 affected DNMT1 and UHRF1 protein. To this aim, we cloned two distinct short-hairpin RNAs (shRNAs) against *Usp7* (referred to as sh1 and sh2), and knockdown of *Usp7* was confirmed by Western blot (**Figure 6K**). Silencing of USP7 suppressed DNMT1 and UHRF1 protein abundances, which were recovered after four hours of treatment with MG132 (**Figure 6L**). We then determined whether LSD1 interacted with USP7. Both WT and LSD1^MUT^ interacted with USP7 (**Figure 6M**), indicating that LSD1 functions as a scaffolding protein to promote the deubiquitination of DNMT1 and UHRF1 independently of its demethylase activity. Overall, our data revealed that demethylase independent function of LSD1 modulates DNMT1 and UHRF1 protein levels, and LSD1 serves as a scaffolding protein to regulate their deubiquitination and stability (**Figure 6N**).

## DISCUSSION

Despite the importance of repressive chromatin marks in maintaining ESC identity and transitions to differentiated fates, we still know very little about the mechanism by which LSD1 operates in pluripotency. In this study, we have employed CRISPR/Cas9-mediated genome editing and multi-layered integrative approaches to characterise the function of LSD1 in mouse ESCs. We propose that the demethylase activity of LSD1 does not play a major role in ESC self-renewal, but it is required for proper differentiation. In ESCs, ablation of LSD1 results in decreased DNMT1 and UHRF1 proteins coupled with global hypomethylation. In this scenario, the catalytic activity of LSD1 is not essential as *Lsd1* KO with reexpression of WT LSD1 or catalytically-dead LSD1 recover the amount of DNMT1 and UHRF1 protein, and DNA methylation levels. Our studies are consistent with a model, as schematically illustrated in **Figure 6N**, that LSD1 acts as a scaffolding protein to recruit USP7, inhibiting DNMT1 and UHRF1 ubiquitylation, and consequently controlling UHRF1 and DNMT1 protein stability. Our studies elucidate for the first time a novel mechanism by which the LSD1-USP7 axis coordinates DNA methylation maintenance in mouse ESCs.

LSD1 is required for early embryogenesis [6, 7]; however, our data show that LSD1 is dispensable for ESC self-renewal. In this study, we have identified more than 3,000 common differentially expressed genes in *Lsd1* KO ESCs which are not related to stemness or differentiation. Specifically, upregulated genes were enriched for biological processes linked to cell proliferation and apoptosis, consistent with the defective proliferation phenotype observed upon LSD1 loss. On the other hand, downregulated genes were related to metabolic processes. Studies have demonstrated that ablation of *Lsd1* results in enhanced oxidative capacity in hepatocarcinoma cells and differentiating myoblast, while the reversed metabolic switch is observed during reprogramming [42, 43]. Here, we reveal that deletion of *Lsd1* leads to reduced glycolytic activity without a substantial increase in oxidative respiration in ESCs. Whether this LSD1-mediated metabolic switch has implications during ESC differentiation needs to be further addressed.

To define the role of LSD1 in cell-fate commitment, we assessed the differentiation potential of *Lsd1* KO ESCs using two independent *in vitro* differentiation methods: i) EB-directed differentiation and ii) gastruloids generation. *Lsd1* KO ESCs formed EBs albeit significantly smaller than WT ESCs. In addition, *Lsd1* KO ESCs were unable to form proper gastruloids. Indeed, EBs and gastruloids depleted of LSD1 showed severely compromised expression of lineage determination markers and failed to silence the pluripotency program.

The histone-demethylase independent activity of LSD1 in regulating gene expression programs have recently begun to unravel. Hence, LSD1 catalytic activity has been reported to be dispensable in promoting tumorigenesis in several cancer models where it seems to play a central role as a scaffold for assembling chromatin modifiers and transcription factor complexes [27, 44, 45]. To investigate the relative contributions of LSD1 enzymatic activity and scaffolding functions in pluripotency, we generated double point mutated (A540E/K662A) LSD1 ESCs (LSD1^MUT^), as a recent study has shown that single mutation (K661A) of human LSD1 (K662A in mouse), which has been widely used as a surrogate of a catalytically inactive LSD1, possesses significant H3K4 demethylase activity on nucleosome substrates [28]. Herein, we demonstrate that reconstitution of *Lsd1* KO ESCs with either WT or a catalytically inactive LSD1 recovers the proliferation defect of ESCs, although mutated LSD1 (unlike the WT form) fails to promote proper differentiation. Ablation of LSD1 led to a global increase of H3K4me1; however, such increase of H3K4me1 in regulatory regions is not sufficient to trigger major gene expression changes. Indeed, other studies have found that this histone mark has only a minor effect on the maintenance of enhancer activity and function [46–48], but that enhancers acquiring H3K4me1 are primed for transcription activation upon differentiation [49]. Thus, the retention of H3K4me1 at pluripotency enhancers in *Lsd1* KO ESCs could explain why pluripotency genes do not undergo silencing upon EB and gastruloid formation, in agreement with previous observations [16]. However, our findings substantially differ from the aforementioned study as we also observed LSD1 occupancy at a subset of genes that are induced during neuronal differentiation, experiencing these genes an increase of H3K4me1 deposition upon LSD1 loss. What is the role of LSD1-dependent H3K4me1 demethylation in these neural markers? One possibility would be that those enhancers are also poised in ESCs and might be subjected to a context-dependent expression pattern, wherein they undergo activation under neuronal differentiation and repression when differentiating onto other lineage programs. Nevertheless, whether H3K4me1 has a regulatory function within enhancers or is simply a useful mark to identify them remains unclear [50].

In addition, we found that knockout of *Lsd1* can elicit genome-wide loss of DNA methylation. In mammals, DNA and histone methylation are highly interrelated to maintain cellular epigenomic landscapes [51]. Hence, repressive histone marks, such as H3K9me3, usually coexist with DNA methylation to generate local formation of heterochromatin. Additionally, there is a strong anti-correlation between CpG methylation and H3K4 methylation that is particularly pronounced at CpG islands [52]. Particularly in our research context, it has been shown that LSD1-mediated demethylation of H3K4me1 is critical in guiding de novo DNA methylation at enhancers of pluripotency genes [23]. We speculate that some degree of DNA hypomethylation in *Lsd1* KO ESCs may be attributed to decreased H3K9me2/3 and increased H3K4me1 levels that exhibit these cells.

LSD1-mediated demethylation of DNMT1 and UHRF1, which has been reported to impede their proteasome degradation [6, 20], has been proposed as an alternative mechanism underlying the molecular crosstalk between DNA and histone methylation. However, our evidence strongly indicates that both LSD1^WT^ and LSD1^MUT^, although to a less extent, are able to recover DNA methylation levels by two non-mutually exclusive mechanisms: i) maintaining DNMT1 and UHRF1 protein stability and ii) recruiting DNMT1 at specific genomic loci, suggesting that the demethylase activity of LSD1 is dispensable to maintain DNA methylation in ESCs. In line with this, targeting LSD1 with the catalytic-specific irreversible inhibitor GSK_LSD1 had no impact on DNMT1 and UHRF1 protein levels, and global DNA methylation. Recently, UM171, a pyrimidoindole derivative, has been found to promote the degradation of the LSD1-containing chromatin remodeling complex CoREST in hematopoietic stem cells [53]. This compound provides a potential means to investigate how LSD1 functions as a scaffold in chromatin dynamics. Lysine methylation can modulate numerous molecular processes by fine-tuning non-histone protein function [54]. Amongst them, lysine methylation has been shown to regulate protein stability. Hence, several studies have shown that lysine methylation can either block polyubiquitination-dependent proteasomal degradation or promote ubiquitination by recruiting the ubiquitin ligase machinery. Even though emerging evidence has revealed that LSD1 can demethylate non-histone proteins [8], structural biology studies show that there is insufficient space in the catalytic center of LSD1 to accommodate more than three residues N-terminal to the target lysine residue [28]. Albeit, we cannot rule out the possibility that a lysine-methylation-dependent proteolytic mechanism controls DNMT1 and UHRF1 protein stability, we propose that LSD1 facilitates deubiquitylation independent of its demethylase activity in ESCs as both WT and LSD1^MUT^ are able to interact with USP7. Whether DNMT1 and UHRF1 methylation status is required for LSD1 recognition needs to be further addressed. A similar example can be found in the serine/threonine kinase AKT by which AKT K64 methylation is required to trigger ubiquitination of AKT following growth factor stimulation. The interaction between AKT and the E3 ligases is mediated through the histone demethylase JMJD2A, which recognizes methylated AKT independent of its catalytic activity [55].

In summary, we demonstrate that both catalysis of histone demethylation and scaffolding function of LSD1 may be variably important for control gene expression depending on the cellular context i.e., in pluripotency or during ESC differentiation. Our results prompt a re-evaluation of the proposed mechanism of action for LSD1 in demethylating non-histone substrates, especially DNMT1 and UHRF1, to increase their stability. They also bring light to a new LSD1-USP7 axis to coordinate DNA methylation maintenance in mouse ESCs.

### Limitations of the study

The proteomics analysis indicated both stabilizing and destabilizing effects of LSD1 on the global protein expression profile. However, the specific role of LSD1 in destabilization was not further analyzed in our study. Understanding the mechanisms by which LSD1 mediates protein destabilization would provide valuable information on the broader regulatory functions of LSD1 in pluripotency. Secondly, in addition to USP7-mediated regulation, we cannot rule out the possibility that other protein deubiquitinases are involved in promoting the LSD1-mediated stability of DNMT1 and UHRF1. Finally, although we propose that DNMT1 and UHRF1 are not substrates of LSD1, we cannot rule out the possibility that DNMT1 and UHRF1 need to be methylated in order to be recognized by LSD1 and guide them for deubiquitination.

### Experimental procedures

#### Antibodies

The following commercially available antibodies were used for western blot: anti-LSD1 (Abcam, ab17721, 1:2500), anti-DNMT1 (Abcam, ab188453, 1:2500), anti-UHRF1 (Invitrogen, PA5-29884, 1:2000), anti-DNMT3A (Abcam, ab188470, 1:2500), anti-DNMT3B (Abcam, ab79822, 1:2000), anti-DNMT3L (Abcam, ab194094, 1:2000), anti-OCT4 (Santa-Cruz, SC-8628, 1:1500), anti-MYC (Cell Signalling, 2276S, 1:4000), anti-βACTIN (Sigma, A54411:5000), anti-H3 (Abcam, ab8895, 1:8000), anti-USP7 (Invitrogen, PA5-34911, 1:2000)). ChIP was performed with anti-H3K4me1 (Abcam, ab8895, 2ug), anti-LSD1 (Abcam, ab17721, 10ug), and anti-DNMT1 (Abcam, ab19905, 10ug). Co-IP experiments were performed with the following antibodies: LSD1 (Abcam, ab17721, 5ug) and Rabbit IgG (Abcam, ab37415, 5ug). Dot-blot experiments were performed with anti-5hmC (Active Motif, 39791, 1:4000) and anti-5mC (Active Motif, 39649, 1:4000). For IF staining we used anti-LSD1 (Abcam, ab129195, 1:500), anti-5hmC (Active Motif, 39791, 1:250), anti-5mC (Active Motif, 39649 1:250), anti-OCT4 (Santa-Cruz, SC-8628, 1:250), anti-SSEA1 (Thermo Scientific, MA5-17042, 1:250), anti-Rabbit (Thermo Scientific, A-11011, 1:1000), and anti-Mouse (Thermo Scientific, A11029, 1:1000).

#### Constructs

To generate the rescue construct, the cDNA of *Lsd1* isoform 2 from mouse ESCs was amplified with RevertAid® First Strand cDNA Synthesis Kit and cloned into a pGEM®-T easy vector (Promega), and ultimately subcloned into the pSIN-E2F_MYC (Sigma-Aldrich) vector using EcoRI as restriction enzymes. All primers used for cloning purposes are described in **Table S9.** A double point mutated (A540E/K662A) mouse LSD1 construct was generated using QuikChange Multi Site-Directed Mutagenesis Kit following the manufacturer’s protocol.

#### Cell Culture

CCE murine ESCs were maintained on 0.1% gelatin-coated tissue culture plates under feeder-free culture conditions. The complete media composition consists of Dulbecco’s modified Eagle’s medium (DMEM) high glucose (Gibco, 41966-029), 15% fetal bovine serum (FBS; Gibco, 10500-064), 1% MEM non-essential amino acids (Sigma-Aldrich, M7145), 0.1 mM of 2-β mercaptoethanol, 1% l-glutamine (Hyclone, SH30034.01) and 1% penicillin/streptomycin (Gibco, 15140-122). Complete media was supplemented with Leukemia inhibitory factor (LIF; R&D systems, 8878-LIF-100/CF, 0.01 ng/µL). All cell cultures were maintained at 37°C with 5% CO_2_.

### RA and EB differentiation assays

To induce retinoic acid (RA)-mediated differentiation, 2.1 × 10^3^ cells/cm^2^ of mouse ESCs were seeded onto tissue culture plates pre-coated with 0.1% gelatin in complete media without LIF and with 5 μM RA (Sigma-Aldrich).

For EB differentiation, ESCs at a density of 8.8 × 10^4^ cells/cm^2^ were seeded in low attachment plates in the presence of a complete medium without LIF. The medium was replenished every 48 h. EB sizes were quantified using NIS-elements software.

### Generation of CRISPR/Cas9 Knockouts

ESCs were seeded in a 12 wells-plate and transfected with 0.8 µg of Cas9 expression vector PX459 (Addgene plasmid #62988) containing the corresponding sgRNAs using Lipofectamine (Invitrogen). All sgRNA-Cas9 plasmids were obtained by ligation (T7 DNA ligase, Fermentas) of annealed complementary oligonucleotides with the PX459 vector (digested with BbsI (BpilI) (Thermo Scientific). All sgRNAs were designed using the Zhang lab’s online tool. Knockouts (KOs) were screened by PCR and validated by western blot and sanger sequencing. The specific primers flanking the cleavage sites were designed to detect the insertion/deletion (INDEL) in the target regions (**Table S9**).

### Mismatch Detection Assay by T7 Endonuclease of *Lsd1* knockouts

Genomic DNA was extracted using the GeneJET Genomic DNA Purification kit (Thermo Scientific, K0722) following the manufacturer’s instructions. Target regions were PCR-amplified (**Table S9**) with DreamTaq 2X Master Mix (Thermo Scientific, K1081). PCR products were denatured at 95 °C for 2 min and re-annealed at -2°C /second temperature ramp to 85°C, followed by a -0.1°C /second temperature ramp to 25°C. The PCR products (20 µL) were incubated with 5 U T7E1 enzyme (NEB #E3321) at 37°C for 20 min. Products from mismatch assays were separated by electrophoresis on a 2% agarose gel in TAE.

### Alkaline phosphatase activity

Alkaline phosphatase (AP) activity was measured using the Stemgent Alkaline Phosphatase Staining kit (Stemgent), following the manufacturer’s recommendations.

### Immunofluorescence staining

For immunofluorescence staining, cells were fixed in 4% paraformaldehyde (PFA) at room temperature (RT) for 15 min, followed by permeabilization at RT for 30 min with 0.25% Tween-20 in PBS (PBST). Cells were then washed twice with 1X PBST and blocked at 10% in normal goat serum (Invitrogen), 1% bovine albumin serum (Hyclone), and 0.05% Tween 20) at RT for 1 hour. The cells were stained with the specified primary antibodies overnight at 4°C in a blocking buffer followed by secondary antibodies staining at RT for 1 h. 6-diamino-2-phenylindole (DAPI) was used for nuclear DNA staining (4 min). Images were acquired using a Zeiss microscope.

### RT-qPCR analysis

Total RNA was extracted using the RNeasy Mini Kit (Qiagen). 1 μg of total RNA was reverse transcribed using the RevertAid First Strand cDNA Synthesis kit (Invitrogen). Quantitative PCR (qPCR) was performed using the Power Up SYBR Green qPCR Master Mix (Applied Biosystems). Gene expression-specific primers used for this study are listed in **Table S9**.

### RNA-Seq Library Preparation

RNA-Seq library preparation was performed at Novogene (Hong Kong). Samples were sequenced by the Illumina HiSeqTM platform (Illumina) as 100 bp pair-ended reads.

### RNA-Seq Analysis

Contaminant (aligned) RNA-Seq reads were filtered and aligned to several mouse reference databases including the mouse genome (mm9, NCBI Build 37). Differentially expressed genes were identified by the R package DEGseq [56] using a false discovery rate (FDR) < 0.001 and fold-change > 1.5.

### Cellular Proliferation

2 x 10^4^ cells were plated in 6-well plate and counted every second day using trypan blue (BioRad).

### Apoptosis Assay, and Cell Cycle Analyses

Apoptosis assay, and cell cycle analysis were performed using Muse™ Cell Analyzer from Millipore following the manufacturer’s instructions.

### Metabolic Analysis

Glycolysis and oxygen phosphorylation rate were measured using Agilent Seahorse XF Glycolysis Stress Test Kit and Agilent Seahorse XF Cell Mito Stress Test Kit by evaluating extracellular acidification rate and oxygen consumption rate, respectively, according to the manufacturer’s instructions.

### Gastruloids aggregation assay

Gastruloids were generated following the previously described protocol [29, 57]. In brief, 300 ESCs were seeded onto low attachment u-bottomed 96-well plates in 40 μl N2B27 media (50% Neurobasal medium, Gibco), 50% advanced DMEM (Gibco), 1x N2 (Gibco, 17502048), 1x B27 (Gibco, 7504044), 2 mM glutamine (Hyclone, SH30034.01), 1x penicillin/streptomycin (Gibco, 15140-122), and 0.1 mM of 2-β mercaptoethanol. After 48 h of incubation, 3 μM CHIR (4423, Tocris Biosciences) was added for 12 h and replenished with N2B27 media. Gastruloids were harvested for total RNA and total protein after 120 h of aggregate formation. Media was changed every day and cells were maintained at 37°C with 5% CO2. The gastruloids length was quantified using NIS-elements software.

### Whole cell extract preparation

Cells were washed with cold 1X PBS, pelleted, and incubated with lysis buffer containing 50 mM Hepes pH 7.5, 150 mM NaCl, 3 mM MgCl2, 0,2% Triton X-100 0.2% Nonidet NP-40, 10% glycerol supplemented with a protease inhibitor in ice for 15 mins. Then, lysates were sonicated for ten cycles with 30 s pulses on/off and centrifuged at 13,000 rpm for 15 mins. The supernatant containing the whole cell extract was frozen at -80°C for further analysis.

### Nuclear extraction

Cells were washed with cold PBS, scraped off, and pelleted. The pellet was resuspended in at least 5 volumes of buffer A (10 mM HEPES pH 7.9, 1.5 mM MgCl_2_, 10 mM KCl, 2 mM DTT) in the presence of protease inhibitors (Fisher Scientific) and incubated for 10 min on ice. After centrifugation, the pellet was resuspended in 2 volumes of buffer A, douncer-homogenized 10 times, and centrifuged at maximum speed for 10 min. Nuclei pellets were then resuspended in 2 volumes of buffer B (20 mM HEPES pH 7.9, 1.5 mM MgCl_2_, 500 mM NaCl, 25% Glycerol, 0.5 mM EDTA, 1 mM DTT) supplemented with protease inhibitors, incubated on a rotator at 4°C for at least 30 min, and at maximum speed for 20 min. The supernatant containing the nuclear fraction was frozen at -80°C for further analysis.

### Subcellular Protein fractionation

The cytosolic, soluble nuclear, and chromatin-bound proteins were separated following the manufacturer’s protocol of the Subcellular protein fractionation kit (Thermo Scientific).

### Co-immunoprecipitation and Immunoblotting

For immunoprecipitation experiments, 1 mg of nuclear extracts were pre-cleared with protein A magnetic beads (BioRad) for 1 hour at 4°C and incubated overnight on a rotator with specific antibodies at 4°C. Following this, protein A magnetic beads were added for 3 h before washing 4 times in ice-cold IP buffer (10 mM Tris; adjust to pH 7.4, 1 mM EDTA, 1 mM EGTA; pH 8.0, 150 mM NaCl, 1% Triton X-100, 0.2 mM sodium orthovanadate) supplemented with protease inhibitors. Immunoprecipitated complexes were resolved by SDS-PAGE, transferred to nitrocellulose membranes (Invitrogen), and immunoblotted with the indicated antibodies, followed by ECL detection (Thermo Scientific).

### DNA Dot Blots

Indicated amounts of genomic DNA were denatured in denaturing buffer (200 mM NaOH, 20 mM EDTA) at 95°C for 5 min and neutralised with an equal volume of neutralising agent (2 M NH_4_CH_3_CO_2_). Nitrocellulose membrane (Thermofisher Scientific) was hydrated in 6X SSC and then “sandwiched” in a Minifold 1 Filtration Manifold (GE Healthcare). Each well was equilibrated with 200 μl of 10x SSC and flushed by gentle suction vacuum twice. The membrane was UV-crosslinked using the default settings. After blocking for 1 hour with 5% milk in PBST, the membrane was incubated with 5mC or 5hmC primary antibodies overnight at 4°C, washed four times in PBST, and incubated with the secondary anti-rabbit antibody for 1 h at RT. The membrane was washed again in 1x PBST and the detection was done using the enhanced chemiluminescence substrate (ECL, Thermo Scientific). DNA levels were normalised with methylene blue staining.

### Whole-genome bisulfite sequencing analysis

Whole-genome bisulfite was performed at Novogene. Paired-end reads from Illumina were aligned to the mouse (mm10) genome using the BWA-meth algorithm [58], and the resulting BAM reads were processed for methylation calling using MethylDackel. We extracted subsets of regions from the final bedgraph methylation file with bedtools [59], as well as other BED files obtained from the UCSC Genome Browser (e.g., SINE, LINE, enhancers, and promoters) or manually curated. Statistical analyses and visualization were performed using MATLAB.

### Quantitative analysis of DNA methylation levels using LC-MS/MS

Quantification of 5mC and 5hmC was performed according to the previously described protocol [60]. Genomic DNA was extracted using the GeneJet DNA purification Kit (Thermofisher Scientific). In brief, extracted DNA was desalted with vertical ultrafiltration and digested by DNase I (New England Biolabs, Ipswich, MA, U.S.A.), 0.02 U snake venom phosphodiesterase (SVP; Worthington Biochemical Corporation, Lakewood, CO, U.S.A.), and 5.0 U calf intestine alkaline phosphatase (CIP; New England Biolabs, Ipswich, MA, U.S.A.) and incubated at 37°C overnight. The digested DNA solutions were filtered by ultrafiltration tubes and then subjected to LC-MS/MS analysis for detection of 5mC and 5hmC.

### Mouse DNA methylation beadChip Array

Samples were bisulfite converted using EZ DNA Methylation-Gold™ Kit (Zymo Research, CA, USA) following the manufacturer’s protocol with modifications for Illumina Infinium Methylation Assay. Infinium Mouse Methylation BeadChip (Illumina, Inc., San Diego, CA, USA) arrays were used to profile genome-wide DNA methylation. This platform interrogates over 285,000 methylation sites per sample at single-nucleotide resolution [61]. Samples were hybridized in the array following the manufacturer’s instructions.

Raw signal intensities were preprocessed and analysed using SeSAMe (v1.14.2) [62]. DNA methylation beta values were obtained from raw IDAT files using the openSesame pipeline within R statistical environment (v4.0.3). Briefly, this pipeline includes a data preprocessing procedure consisting of strain-specific masking for mouse methylation array probes, followed by masking of non-uniquely mapped probes (poor design), channel inference for Infinium I probes, non-linear dye bias correction, detection p-value masking (p > 0.05) using the pOOBAH algorithm and background subtraction using the noob method [35].

Differential methylation loci were computed using DML function in SeSAMe by modelling DNA methylation values (beta values) using mixed linear models. Loci with a False Discovery Rate (FDR) adjusted p-value < 0.05, and absolute beta value difference between conditions > 0.66 were considered significant. Principal component analysis and the corresponding plot were computed using PCAtools (v.2.8.0) R package. Euclidean distance scores and Ward’s minimum variance method were applied to attain hierarchical clustering represented as a heatmap using heatmap.2 function from the gplots (v3.1.3) package in R. Density plots were performed with minfi (v1.42.0) R package, and correlation analyses were computed using Spearman correlation coefficient and plotted using ggplot2 (v3.3.6) package in R. Finally, all downstream analyses were performed within the R statistical environment (v4.0.3).

The DNA methylation analysis was performed using the mm10 mouse genome reference build annotation from the Infinium Mouse Methylation BeadChip Array manifest file (http://zwdzwd.github.io/InfiniumAnnotation#mouse) [35].

### Quantitative analysis of histone modifications using LC-MS/MS

After homogenization of the cells with nuclear extraction buffer (15 mM Tris, 60 mM KCl, 15 mM NaCl, 5 mM MgCl_2_, 1 mM CaCl_2_, and 250 mM sucrose), histone proteins were extracted as previously described [63]. Briefly, histones were acid extracted from nuclei with 0.2 M H_2_SO_4_ for 2 h and precipitated with 33% trichloroacetic acid (TCA) overnight. The pellets, containing histone proteins, were dissolved in 30 μL of 50 mM NH_4_HCO_3_, pH 8.0. Derivatization reagent was prepared by mixing propionic anhydride with acetonitrile in a ratio of 1:3 (v/v), and the reagent was mixed with the histone sample in a ratio of 1:4 (v/v) for 20 min at RT. The reaction was performed twice to ensure derivatization completion. Histones were then digested with trypsin (enzyme: sample ratio 1:20, 6 h, RT) in 50 mM NH_4_HCO_3_. The derivatization procedure was repeated after digestion to derivatize peptide N-termini.

Prior to mass spectrometry analysis, samples were desalted using a 96-well plate filter (Orochem) packed with 1 mg of Oasis HLB C-18 resin (Waters). Briefly, the samples were resuspended in 100 µl of 0.1% TFA and loaded onto the HLB resin, which was previously equilibrated using 100 µl of the same buffer. After washing with 100 µl of 0.1% TFA, the samples were eluted with a buffer containing 70 µl of 60% acetonitrile and 0.1% TFA and then dried in a vacuum centrifuge. Thereafter, Samples were resuspended in 10 µl of 0.1% TFA and loaded onto a Dionex RSLC Ultimate 300 (Thermo Scientific), coupled online with an Orbitrap Fusion Lumos (Thermo Scientific). Chromatographic separation was performed with a two-column system, consisting of a C-18 trap cartridge (300 µm ID, 5 mm length) and a picofrit analytical column (75 µm ID, 25 cm length) packed in-house with reversed-phase Repro-Sil Pur C18-AQ 3 µm resin. Peptides were separated using a 30 min gradient from 1-30% buffer B (buffer A: 0.1% formic acid, buffer B: 80% acetonitrile + 0.1% formic acid) at a flow rate of 300 nl/min. The mass spectrometer was set to acquire spectra in a data-independent acquisition (DIA) mode. Briefly, the full MS scan was set to 300-1100 m/z in the orbitrap with a resolution of 120,000 (at 200 m/z) and an AGC target of 5x10e5. MS/MS was performed in the orbitrap with sequential isolation windows of 50 m/z with an AGC target of 2x10e5 and an HCD collision energy of 30.

Histone peptides raw files were imported into EpiProfile 2.0 software [64]. From the extracted ion chromatogram, the area under the curve was obtained and used to estimate the abundance of each peptide. In order to achieve the relative abundance of post-translational modifications (PTMs), the sum of all different modified forms of a histone peptide was considered as 100% and the area of the particular peptide was divided by the total area for that histone peptide in all of its modified forms. The relative ratio of two isobaric forms was estimated by averaging the ratio for each fragment ion with different mass between the two species. The resulting peptide lists generated by EpiProfile were exported to Microsoft Excel and further processed for a detailed analysis.

### Chromatin immunoprecipitation (ChIP)

Mouse ESCs were chemically crosslinked by adding 1/10 volume of fresh 11% formaldehyde solution for 10 min at RT and quenched by adding 1/20 volume of 2.5 M of glycine for 5 min at RT. Cells were then washed twice with ice-cold PBS, scraped and collected by centrifugation. The cell pellet was first lysed in the lysis buffer 1 (50 mM, Hepes-KOH pH 7.5, 140 mM NaCl, 1 mM EDTA, glycerol (10% vol/vol) NP-40 (0.5% vol/vol), Triton X-100 (0.25% vol/vol)) for 10 min at 4°C, followed by lysis in buffer 2 (200 mM NaCl, 1 mM EDTA, 0.5 mM EGTA, 10mM Tris (pH 8)) with complete protease inhibitors. After lysis, cells were sonicated in lysis buffer 3 (100 mM NaCl, 1 mM EDTA, 0.5 mM EGTA, 10 mM Tris pH 8, Na-Deoxycholate (DOC) (0.1% vol/vol), N-lauroyl sarcosine (0.5% vol/vol) for 16 x 30-second pulses (30-second pause between pulses) at high power in a Bioruptor® Sonication System (Diagenode) and centrifuged for high speed for 15 min. The supernatant containing the chromatin fraction was subjected to immunoprecipitation.

Spike-in control (human MCF10A cell line) was used for normalization of the ChIP-seq reads. For this, human MCF10A chromatin was prepared, as mentioned above. In the ESCs chromatin and antibodies immunoprecipitation complex, 5% of MCF10A (vol/vol) chromatin was added. This complex was mixed with 50 μl of Dynabeads® Protein G magnetic beads and incubated overnight at 4°C. Protein-DNA bead complexes were washed first with RIPA buffer (50 mM Hepes pH 7.6, 1 mM EDTA, DOC (0.7% vol/vol)), NP-40 (1% vol/vol), 0.5 M LiCl) for 5 times and then, with TE containing 50 mM NaCl. The protein-DNA complexes were eluted from the beads twice by incubation with 100 μL of elution buffer (50 mM Tris pH 8, 10 mM EDTA, SDS (1% vol/vol) at 65°C for 15 min with shaking. Reverse crosslinking was performed by adding 200 mM NaCl in the eluate and incubating it at 65°C overnight with shaking. Thereafter, RNA and protein were removed from the samples by treating with RNase A and proteinase K as the manufacturer’s recommendations. DNA was subsequently purified by phenol-chloroform extraction, followed by ethanol precipitation The input sample was also treated for crosslink reversal and following steps after this.

### ChIP-seq analysis

Library construction and paired-end read sequencing (20M per sample) of triplicate immunoprecipitated chromatin and control inputs were performed using Illumina technology at Novogene UK. Raw reads were aligned to the human (hg38) and mouse (mm10) genomes using BWA [58]. The final BAM files were normalized using the number of reads that uniquely mapped to the human genome [65]. Peaks were called using MACS2 [66] with pooled IPs and inputs (FDR < 0.05), and they were annotated using the chIPseekeR [67] package in R. Subsequent analyses were performed using bedtools [59] for peak overlapping, HOMER [68] for motif detection, and MATLAB, GBiB [69], and EaSEQ [70] for visualization.

### ReChIP

Mouse ESCs were chemically crosslinked by the addition 1/10 volume of fresh 11% formaldehyde solution for 10 min at RT and quenched by adding 1/20 volume of 2.5 M of glycine for 5 min at RT. Cells were then washed twice with ice-cold PBS, scraped, and collected by centrifugation. The cell pellet was first lysed in the swelling buffer (25 mM, Hepes-KOH pH 7.9, 1.5 mM MgCl_2_, 10 mM KCl, NP-40 (0.1% vol/vol), 1mM DTT) supplemented with complete protease inhibitors for 10 min at 4°C and homogenized with douncer (20 times up and down). Following centrifugation, the pellet was sonicated in a sonication buffer (50 mM Hepes pH 7.9, 140 mM NaCl, 1 M EDTA, triton X-100 (1% vol/vol), Na-Deoxycholate (0.1% vol/vol), 0.1% SDS, complete protease inhibitors) for 16 x 30-second pulses (30-second pause between pulses) at high power in a Bioruptor® Sonication System (Diagenode) and centrifuged for high speed for 15 min. The recovered mouse chromatin fraction was mixed with 50 μl of Dynabeads® Protein G magnetic beads and incubated with the indicated antibodies overnight at 4°C. Protein-DNA bead complexes were subsequently washed with different buffers for 10 min twice: i) sonication buffer, ii) wash buffer A (50 mM Hepes pH 7.9, 500 mM NaCl, 1 M EDTA, triton X-100 (1% vol/vol), Na-Deoxycholate (DOC) (0.1% vol/vol), 0.1% SDS), iii) wash buffer B (20 mM Tris, pH 8, 1 mM EDTA, 250 mM LiCl, NP-40 (0.5% vol/vol), Na-Deoxycholate (0.5% vol/vol)) and iv) TE buffer. The protein-DNA complexes were divided into two fractions. The first fraction was eluted from the beads twice by incubation with 100 μL of elution buffer (50 mM Tris pH 8, 10 mM EDTA, SDS (1% vol/vol) at 65°C for 15 min with shaking. The second fraction was resuspended with an equal volume of 10 mM DTT for 30 min at 37°C. After elution, fractions were pooled, and 10% of the elute was kept as input for ReChIP with a second antibody. Then, elute was diluted 10 times with sonication buffer, mixed with 50 μl of Dynabeads® Protein G magnetic beads, and incubated with the second antibodies overnight at 4°C. A fraction of elute was mixed with empty beads as an antibody leakage control for the first ChIP. After overnight incubation, protein-DNA complexes were washed, as mentioned above and eluted from the beads twice by incubation with 100 μL of elution buffer (50 mM Tris pH 8, 10 mM EDTA, SDS (1% vol/vol) at 65°C for 15 min with shaking. Reverse crosslinking was performed by adding 200 mM NaCl in the eluate and incubating it at 65°C overnight with shaking. Thereafter, RNA and protein were removed from the samples by treating them with RNase A and proteinase K as per the manufacturer’s recommendations. DNA was subsequently purified by phenol-chloroform extraction, followed by ethanol precipitation. The input sample was also treated for crosslink reversal and following steps after this. Fold enrichment over 10% input was calculated using the 2Delta Ct method. The primer sets used for ChIP analysis are listed in **Table S9**.

### Gene Ontology (GO) Analysis

Gene ontology (GO) analysis was performed using the web tool The Database for Annotation, Visualization and Integrated Discovery (DAVID) (http://david.abcc.ncifcrf.gov/).

### Heatmaps

Heat maps were generated from the log2-transformed expression level for a given gene in a specific sample using Heatmapper.

### His10-SUMO-LSD1 expression and purification

Mouse His10-SUMO-LSD1 constructs were transformed into Rosetta (DE3) and the proteins were expressed by auto-induction media at 20°C overnight. Cells were lysed, and proteins were purified using NiNTA affinity resin (Thermo Scientific, 88222). Briefly, cells were resuspended in lysis buffer (50 mM NaP 8.0, 500 mM NaCl, 10% glycerol, 0.2% Triton X100, 10 mM Imidazole, 1 mM βME, DNAse) and sonicated (15 cycles 10 on/off). Following centrifugation, the lysate was incubated with NiNTA resin at 4°C for 1 h. The resin was washed with 25CV wash buffer 1 (50 mM NaP 8.0, 500 mM NaCl, 10% glycerol, 20 mM Imidazole,1 mM βME) followed by 25CV high salt buffer (50 mM NaP 8.0, 1 M NaCl, 10%glycerol, 20 mM Imidazole, 1 mM βME) and finally with 25CV wash buffer 1. The protein was eluted with buffer containing 50 mM NaP 8.0, 300 mM NaCl, 10% glycerol, 250 mM Imidazole, and 1 mM βME.

The His10SUMO-tag was cleaved off using SUMOprotease 1/100 (w/w) in reaction buffer 20 mM NaP 8.0, 150 mM NaCl, 10%glycerol, 1 mM βME, at 4°C overnight. The tag and protease were removed by NiNTA affinity purification. The protein was further purified by gel filtration on a Superdex200 column (Cytiva) equilibrated in a 20 mM NaP 8.0, 500 mM NaCl, 5% glycerol, and 1 mM DTT buffer.

### Enzymatic activity of LSD1

The enzymatic activity of LSD1 was determined using a peroxidase-coupled assay. Briefly, 0.3 µM of LSD1 was mixed with the reaction buffer containing 50 mM HEPES pH 8.5, 0.1 mM Amplex Red and 0.3 mM horseradish peroxidase. Then, the reaction mixture was added to serially diluted H3K4me2 peptide (Chinapeptides) from 40 µM to 0.31 µM and incubated at RT for 10 min. The fluorescence signal obtained from the conversion of amplex red to resorufin was measured on a Clariostar plate reader (BMG Labtech) with excitation at 510 nm and emission at 595 nm. The intensity of the fluorescence signal is directly proportional to the demethylase activity. Non-linear regression analysis was performed to calculate K_cat_ and K_m_ of enzyme activity.

### LSD1 inhibitors treatment

Mouse ESCs were seeded at a density of 8x10^4^ in a 10 cm plate. After 24 h, cells were treated with a final concentration of 10µM GSK_LSD1. Cells were harvested for whole cell extracts, chromatin, and RNA following 24 h of treatment.

### CHX treatment and proteasome inhibition

Mouse ESCs were seeded in a 6-well plate at the density of 2 × 10^5^ cells and after 24 h, cells were treated with CHX at a final concentration of 50 μg/ml for 0, 3, 6, and 9 h, respectively. For proteasome inhibition, cells were incubated with MG-132 at a final concentration of 20 μg/ml for 4 h. Cells were harvested for whole cell extracts on indicated time points, and proteins were subjected to immunoblotting. Blots were quantified using Image Lab software.

### Mass spectrometry-based proteomics

5 μg of whole cell extracts were denatured with a final concentration of 2% SDS and 20 mM TCEP. Samples were digested with a modified sp3 protocol, as previously described [71, 72]. Briefly, samples were added to a bead suspension (10 μg of beads (Sera-Mag Speed Beads, 4515-2105-050250, 6515-2105-050250) in 10 μl 15% formic acid and 30 μl ethanol) and incubated shaking for 15 min at room temperature. Beads were then washed four times with 70% ethanol. Proteins were digested overnight by adding 40 μl of 5 mM chloroacetamide, 1.25 mM TCEP, and 200 ng trypsin in 100 mM HEPES pH 8.5. Peptides were eluted from the beads and dried under a vacuum. Peptides were then labelled with TMTpro (Thermo Fisher Scientific), pooled and desalted with solid-phase extraction using a Waters OASIS HLB Elution Plate (30 μm). Samples were fractionated onto 48 fractions on a reversed-phase C18 system running under high pH conditions, pooling every twelve fractions together. Samples were analyzed by LC-MS/MS using a data-dependent acquisition strategy on a Thermo Fisher Scientific Vanquish Neo LC coupled with a Thermo Fisher Scientific Orbitrap Exploris 480. Raw files were processed with MSFragger [73] against a *Mus musculus* FASTA database downloaded from UniProt (UP000000589) using standard settings for TMT. Data were normalized using vsn [74], and statistical significance was determined using limma [75].

### Statistical Analysis

All values were expressed as mean ± SD. Statistical analysis was performed by the Unpaired Student’s t-test, One-way ANOVA and 2way ANOVA. A probability value of p < 0.05 was considered statistically significant.

### Data availability

All next-generation sequencing data will be publicly accessed in ArrayExpress webserver (pending for accession number). By now, all next-generation sequencing data has been uploaded in https://figshare.com/s/fc83af7bfb83ef37b5d7

The mass spectrometry proteomics data have been deposited to the ProteomeXchange Consortium via the PRIDE partner repository with the dataset identifier PXD042495.

## Acknowledgements

We thank Filipe Pereira (WCMM and Lund University, Sweden) for providing *Dnmt1* and double *Dnmt3a/b* KO ESCs. We would like to thank Aguilo Lab members for useful discussion, the Biochemical Imaging Centre Umeå, and Mikael Lindberg (Protein expertise platform) for assisting with the cloning. This research was supported by grants from the Knut and Alice Wallenberg Foundation, Umeå University, Västerbotten County Council, Swedish Research Council (2017-01636; 2022-01322), Kempe Foundation (JCK-2150), and Cancerfonden (190337 Pj; 22 2455 Pj).

## Author contributions

S.M. and F.A. conceived and designed the study. S.M. and J.G. performed seahorse. J.C. and A.M. performed enzymatic assays of LSD1. S.S. and S.S. performed quantitative analysis of histone modifications using LC-MS/MS. H.W., W.L., and C.L. performed LC-MS/MS on DNA. C.G-P., D.A-E., and M.E. performed methylation studies. A.M. performed the proteomics analysis. A-C.R. performed bioinformatics analysis. S.M., K.K., C.M-G., P.A.S. and F.A. performed experiments. S.M. and F.A. wrote the manuscript. All authors reviewed and edited the manuscript. F.A. supervised the study.

## Supplemental Information

Document S1. Figures S1–S6

Table S1. RNA-seq in WT, *Lsd1* KO1 and *Lsd1* KO2 mouse ESCs.

Table S2. LC-MS/MS of histone modifications in WT, *Lsd1* KO1 and *Lsd1* KO2 mouse ESCs.

Table S3. LSD1 ChIP-seq in WT mouse ESCs.

Table S4. H3K4me1 ChIP-seq in WT and *Lsd1* KO2 mouse ESCs.

Table S5. Differentially methylated positions (DMPs) obtained from the Mouse Methylation MM285 BeadChIP microarray in *Lsd1* KO2, LSD1^WT^ and LSD1^MUT^ ESCs compared to WT.

Table S6. RNA-seq and DMPs correlation analysis betweenWT and *Lsd1* KO2 mouse ESCs.

Table S7. DNMT1 ChIP-seq in WT and *Lsd1* KO2 mouse ESCs.

Table S8: Total proteomics analysis in *Lsd1* KO2, LSD1^WT^ and LSD1^MUT^ ESCs compared to WT.

Table S9. Sequence of primers used in this study.

## References

1. Atlasi, Y. and H.G. Stunnenberg, The interplay of epigenetic marks during stem cell differentiation and development. Nat Rev Genet, 2017. 18(11): p. 643–658.

2. Boland, M.J., K.L. Nazor, and J.F. Loring, Epigenetic regulation of pluripotency and differentiation. Circ Res, 2014. 115(2): p. 311–24.

3. Malla, S., D. Melguizo-Sanchis, and F. Aguilo, Steering pluripotency and differentiation with N(6)-methyladenosine RNA modification. Biochim Biophys Acta Gene Regul Mech, 2019. 1862(3): p. 394–402.

4. Shi, Y., et al., Histone demethylation mediated by the nuclear amine oxidase homolog LSD1. Cell, 2004. 119(7): p. 941–53.

5. Forneris, F., et al., Histone demethylation catalysed by LSD1 is a flavin-dependent oxidative process. FEBS Lett, 2005. 579(10): p. 2203–7.

6. Wang, J., et al., The lysine demethylase LSD1 (KDM1) is required for maintenance of global DNA methylation. Nat Genet, 2009. 41(1): p. 125–9.

7. Wang, J., et al., Opposing LSD1 complexes function in developmental gene activation and repression programmes. Nature, 2007. 446(7138): p. 882–7.

8. Martinez-Gamero, C., S. Malla, and F. Aguilo, LSD1: Expanding Functions in Stem Cells and Differentiation. Cells, 2021. 10(11).

9. Adamo, A., et al., LSD1 regulates the balance between self-renewal and differentiation in human embryonic stem cells. Nat Cell Biol, 2011. 13(6): p. 652–9.

10. Han, X., et al., Destabilizing LSD1 by Jade-2 promotes neurogenesis: an antibraking system in neural development. Mol Cell, 2014. 55(3): p. 482–94.

11. Foster, C.T., et al., Lysine-specific demethylase 1 regulates the embryonic transcriptome and CoREST stability. Mol Cell Biol, 2010. 30(20): p. 4851–63.

12. Choi, J., et al., Histone demethylase LSD1 is required to induce skeletal muscle differentiation by regulating myogenic factors. Biochem Biophys Res Commun, 2010. 401(3): p. 327–32.

13. Musri, M.M., et al., Histone demethylase LSD1 regulates adipogenesis. J Biol Chem, 2010. 285(39): p. 30034–41.

14. Saleque, S., et al., Epigenetic regulation of hematopoietic differentiation by Gfi-1 and Gfi-1b is mediated by the cofactors CoREST and LSD1. Mol Cell, 2007. 27(4): p. 562–72.

15. Su, S.T., et al., Involvement of histone demethylase LSD1 in Blimp-1-mediated gene repression during plasma cell differentiation. Mol Cell Biol, 2009. 29(6): p. 1421–31.

16. Whyte, W.A., et al., Enhancer decommissioning by LSD1 during embryonic stem cell differentiation. Nature, 2012. 482(7384): p. 221–5.

17. Hahm, J.Y., et al., Methylation of UHRF1 by SET7 is essential for DNA double-strand break repair. Nucleic Acids Res, 2019. 47(1): p. 184–196.

18. Majello, B., et al., Expanding the Role of the Histone Lysine-Specific Demethylase LSD1 in Cancer. Cancers (Basel), 2019. 11(3).

19. Mancini, M., et al., The multi-functionality of UHRF1: epigenome maintenance and preservation of genome integrity. Nucleic Acids Res, 2021. 49(11): p. 6053–6068.

20. Zhang, H., et al., SET8 prevents excessive DNA methylation by methylation-mediated degradation of UHRF1 and DNMT1. Nucleic Acids Res, 2019. 47(17): p. 9053–9068.

21. Esteve, P.O., et al., Regulation of DNMT1 stability through SET7-mediated lysine methylation in mammalian cells. Proc Natl Acad Sci U S A, 2009. 106(13): p. 5076–81.

22. Leng, F., et al., Methylated DNMT1 and E2F1 are targeted for proteolysis by L3MBTL3 and CRL4(DCAF5) ubiquitin ligase. Nat Commun, 2018. 9(1): p. 1641.

23. Petell, C.J., et al., An epigenetic switch regulates de novo DNA methylation at a subset of pluripotency gene enhancers during embryonic stem cell differentiation. Nucleic Acids Res, 2016. 44(16): p. 7605–17.

24. Gu, F., et al., Biological roles of LSD1 beyond its demethylase activity. Cell Mol Life Sci, 2020. 77(17): p. 3341–3350.

25. Chao, A., et al., Lysine-specific demethylase 1 (LSD1) destabilizes p62 and inhibits autophagy in gynecologic malignancies. Oncotarget, 2017. 8(43): p. 74434–74450.

26. Lan, H., et al., LSD1 destabilizes FBXW7 and abrogates FBXW7 functions independent of its demethylase activity. Proc Natl Acad Sci U S A, 2019. 116(25): p. 12311–12320.

27. Garcia-Martinez, L., et al., Endocrine resistance and breast cancer plasticity are controlled by CoREST. Nat Struct Mol Biol, 2022. 29(11): p. 1122–1135.

28. Kim, S.A., et al., Crystal Structure of the LSD1/CoREST Histone Demethylase Bound to Its Nucleosome Substrate. Mol Cell, 2020. 78(5): p. 903–914 e4.

29. Beccari, L., et al., Multi-axial self-organization properties of mouse embryonic stem cells into gastruloids. Nature, 2018. 562(7726): p. 272-276.

30. Jung, H.R., et al., Quantitative mass spectrometry of histones H3.2 and H3.3 in Suz12-deficient mouse embryonic stem cells reveals distinct, dynamic post-translational modifications at Lys-27 and Lys-36. Mol Cell Proteomics, 2010. 9(5): p. 838–50.

31. Ortabozkoyun, H., et al., CRISPR and biochemical screens identify MAZ as a cofactor in CTCF-mediated insulation at Hox clusters. Nat Genet, 2022. 54(2): p. 202–212.

32. Guo, G. and A. Smith, A genome-wide screen in EpiSCs identifies Nr5a nuclear receptors as potent inducers of ground state pluripotency. Development, 2010. 137(19): p. 3185–92.

33. Aguilo, F., et al., Coordination of m(6)A mRNA Methylation and Gene Transcription by ZFP217 Regulates Pluripotency and Reprogramming. Cell Stem Cell, 2015. 17(6): p. 689–704.

34. Agarwal, S., et al., KDM1A maintains genome-wide homeostasis of transcriptional enhancers. Genome Res, 2021. 31(2): p. 186–97.

35. Zhou, W., et al., DNA methylation dynamics and dysregulation delineated by high-throughput profiling in the mouse. Cell Genom, 2022. 2(7).

36. Elliott, E.N., K.L. Sheaffer, and K.H. Kaestner, The ’de novo’ DNA methyltransferase Dnmt3b compensates the Dnmt1-deficient intestinal epithelium. Elife, 2016. 5.

37. Mohammad, H.P., et al., A DNA Hypomethylation Signature Predicts Antitumor Activity of LSD1 Inhibitors in SCLC. Cancer Cell, 2015. 28(1): p. 57–69.

38. Carnesecchi, J., et al., ERRalpha induces H3K9 demethylation by LSD1 to promote cell invasion. Proc Natl Acad Sci U S A, 2017. 114(15): p. 3909–3914.

39. Du, Z., et al., DNMT1 stability is regulated by proteins coordinating deubiquitination and acetylation-driven ubiquitination. Sci Signal, 2010. 3(146): p. ra80.

40. Ahmad, T., et al., TIP60 governs the auto-ubiquitination of UHRF1 through USP7 dissociation from the UHRF1/USP7 complex. Int J Oncol, 2021. 59(5).

41. Yi, L., et al., Stabilization of LSD1 by deubiquitinating enzyme USP7 promotes glioblastoma cell tumorigenesis and metastasis through suppression of the p53 signaling pathway. Oncol Rep, 2016. 36(5): p. 2935–2945.

42. Sakamoto, A., et al., Lysine Demethylase LSD1 Coordinates Glycolytic and Mitochondrial Metabolism in Hepatocellular Carcinoma Cells. Cancer Res, 2015. 75(7): p. 1445–56.

43. Sun, H., et al., Lysine-specific histone demethylase 1 inhibition promotes reprogramming by facilitating the expression of exogenous transcriptional factors and metabolic switch. Sci Rep, 2016. 6: p. 30903.

44. Sehrawat, A., et al., LSD1 activates a lethal prostate cancer gene network independently of its demethylase function. Proc Natl Acad Sci U S A, 2018. 115(18): p. E4179–E4188.

45. Hatzi, K., et al., Histone demethylase LSD1 is required for germinal center formation and BCL6-driven lymphomagenesis. Nat Immunol, 2019. 20(1): p. 86–96.

46. Dorighi, K.M., et al., Mll3 and Mll4 Facilitate Enhancer RNA Synthesis and Transcription from Promoters Independently of H3K4 Monomethylation. Mol Cell, 2017. 66(4): p. 568–576 e4.

47. Local, A., et al., Identification of H3K4me1-associated proteins at mammalian enhancers. Nat Genet, 2018. 50(1): p. 73–82.

48. Rickels, R., et al., Histone H3K4 monomethylation catalyzed by Trr and mammalian COMPASS-like proteins at enhancers is dispensable for development and viability. Nat Genet, 2017. 49(11): p. 1647–1653.

49. Wang, C., et al., Enhancer priming by H3K4 methyltransferase MLL4 controls cell fate transition. Proc Natl Acad Sci U S A, 2016. 113(42): p. 11871–11876.

50. Rada-Iglesias, A., Is H3K4me1 at enhancers correlative or causative? Nat Genet, 2018. 50(1): p. 4–5.

51. Li, Y., X. Chen, and C. Lu, The interplay between DNA and histone methylation: molecular mechanisms and disease implications. EMBO Rep, 2021. 22(5): p. e51803.

52. Meissner, A., et al., Genome-scale DNA methylation maps of pluripotent and differentiated cells. Nature, 2008. 454(7205): p. 766–70.

53. Subramaniam, A., et al., Lysine-specific demethylase 1A restricts ex vivo propagation of human HSCs and is a target of UM171. Blood, 2020. 136(19): p. 2151–2161.

54. Cornett, E.M., et al., Lysine Methylation Regulators Moonlighting outside the Epigenome. Mol Cell, 2019. 75(6): p. 1092–1101.

55. Wang, G., et al., SETDB1-mediated methylation of Akt promotes its K63-linked ubiquitination and activation leading to tumorigenesis. Nat Cell Biol, 2019. 21(2): p. 214–225.

56. Wang, L., et al., DEGseq: an R package for identifying differentially expressed genes from RNA-seq data. Bioinformatics, 2010. 26(1): p. 136–8.

57. Baillie-Johnson, P., et al., Generation of Aggregates of Mouse Embryonic Stem Cells that Show Symmetry Breaking, Polarization and Emergent Collective Behaviour In Vitro. J Vis Exp, 2015(105).

58. Li, H. and R. Durbin, Fast and accurate short read alignment with Burrows-Wheeler transform. Bioinformatics, 2009. 25(14): p. 1754–60.

59. Quinlan, A.R. and I.M. Hall, BEDTools: a flexible suite of utilities for comparing genomic features. Bioinformatics, 2010. 26(6): p. 841–2.

60. Lai, W., C. Lyu, and H. Wang, Vertical Ultrafiltration-Facilitated DNA Digestion for Rapid and Sensitive UHPLC-MS/MS Detection of DNA Modifications. Anal Chem, 2018. 90(11): p. 6859–6866.

61. Garcia-Prieto, C.A., et al., Validation of a DNA methylation microarray for 285,000 CpG sites in the mouse genome. Epigenetics, 2022. 17(12): p. 1677–1685.

62. Zhou, W., et al., SeSAMe: reducing artifactual detection of DNA methylation by Infinium BeadChips in genomic deletions. Nucleic Acids Res, 2018. 46(20): p. e123.

63. Karch, K.R., S. Sidoli, and B.A. Garcia, Identification and Quantification of Histone PTMs Using High-Resolution Mass Spectrometry. Methods Enzymol, 2016. 574: p. 3–29.

64. Yuan, Z.F., et al., EpiProfile 2.0: A Computational Platform for Processing Epi-Proteomics Mass Spectrometry Data. J Proteome Res, 2018. 17(7): p. 2533–2541.

65. Danecek, P., et al., Twelve years of SAMtools and BCFtools. Gigascience, 2021. 10(2).

66. Zhang, Y., et al., Model-based analysis of ChIP-Seq (MACS). Genome Biol, 2008. 9(9): p. R137.

67. Yu, G., L.G. Wang, and Q.Y. He, ChIPseeker: an R/Bioconductor package for ChIP peak annotation, comparison and visualization. Bioinformatics, 2015. 31(14): p. 2382–3.

68. Heinz, S., et al., Simple combinations of lineage-determining transcription factors prime cis-regulatory elements required for macrophage and B cell identities. Mol Cell, 2010. 38(4): p. 576–89.

69. Haeussler, M., et al., Navigating protected genomics data with UCSC Genome Browser in a Box. Bioinformatics, 2015. 31(5): p. 764–6.

70. Lerdrup, M., et al., An interactive environment for agile analysis and visualization of ChIP-sequencing data. Nat Struct Mol Biol, 2016. 23(4): p. 349–57.

71. Hughes, C.S., et al., Ultrasensitive proteome analysis using paramagnetic bead technology. Mol Syst Biol, 2014. 10(10): p. 757.

72. Mateus, A., et al., The functional proteome landscape of Escherichia coli. Nature, 2020. 588(7838): p. 473–478.

73. Kong, A.T., et al., MSFragger: ultrafast and comprehensive peptide identification in mass spectrometry-based proteomics. Nat Methods, 2017. 14(5): p. 513–520.

74. Huber, W., et al., Variance stabilization applied to microarray data calibration and to the quantification of differential expression. Bioinformatics, 2002. 18 Suppl 1: p. S96–104.

75. Ritchie, M.E., et al., limma powers differential expression analyses for RNA-sequencing and microarray studies. Nucleic Acids Res, 2015. 43(7): p. e47.

