## Supplementary information for "The catalytic-independent function of LSD1 modulates the epigenetic landscape of mouse embryonic stem cells"

**Supplemental Information**

**Index of supplementary figures:**

**Figure S1.** The ESC transcriptome is not altered upon *Lsd1* loss, related to Figure 1.
**Figure S2.** *Lsd1* deletion halts differentiation, related to Figure 2.
**Figure S3.** Genetic deletion of *Lsd1* affects H3K4me1 and H3K9me2/3, related to Figure 3.
**Figure S4.** Loss of *Lsd1* affects global DNA methylation, related to Figure 4.
**Figure S5.** LSD1 regulates DNMT1 and UHRF1 stability, related to Figure 5.
**Figure S6.** Silencing of *Lsd1* resulted in decreased DNMT1 and UHRF1 stability, related to Figure 6.

**Index of supplementary tables:**

**Table S1 (provided as a spreadsheet):** RNA-seq in WT, *Lsd1* KO1 and *Lsd1* KO2 mouse ESCs.
**Table S2 (provided as a spreadsheet):** LC-MS/MS of histone modifications in WT, *Lsd1* KO1 and *Lsd1* KO2 mouse ESCs.
**Table S3 (provided as a spreadsheet):** LSD1 ChIP-seq in WT mouse ESCs.
**Table S4 (provided as a spreadsheet):** H3K4me1 ChIP-seq in WT and *Lsd1* KO2 mouse ESCs.
**Table S5 (provided as a spreadsheet):** Differentially methylated positions (DMPs) obtained from the Mouse Methylation MM285 BeadChIP microarray in *Lsd1* KO2, $LSD1^{WT}$ and $LSD1^{MUT}$ ESCs compared to WT.
**Table S6 (provided as a spreadsheet):** RNA-seq and DMPs correlation analysis between WT and *Lsd1* KO2 mouse ESCs.
**Table S7 (provided as a spreadsheet):** DNMT1 ChIP-seq in WT and *Lsd1* KO2 mouse ESCs.
**Table S8 (provided as a spreadsheet):** Total proteomics analysis in *Lsd1* KO2, $LSD1^{WT}$ and $LSD1^{MUT}$ ESCs compared to WT.
**Table S9 (provided as a spreadsheet):** Sequence of primers used in this study.

**Figure S1.**

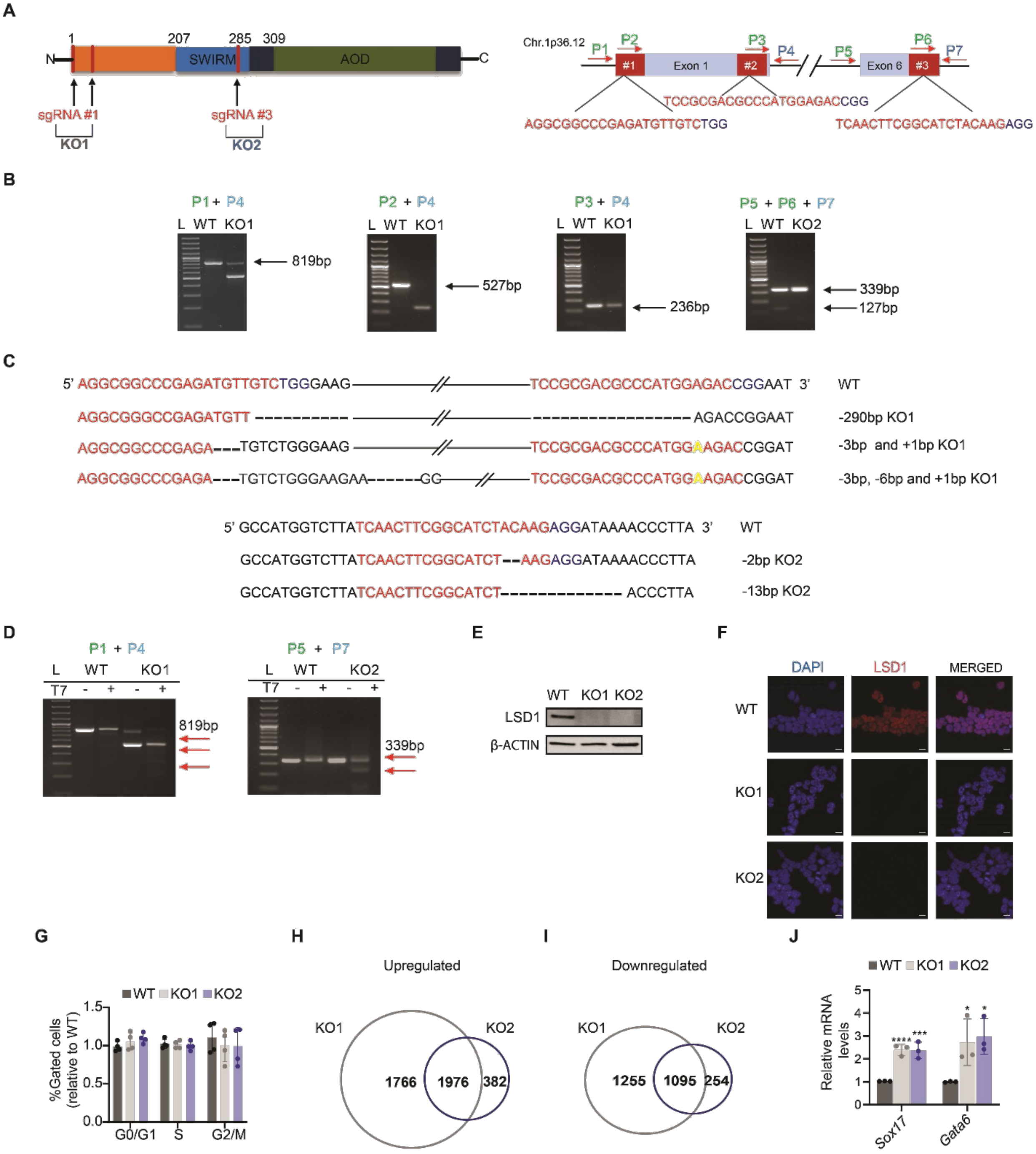


*Figure legends in the next page.*

**Figure S1. The ESC transcriptome is not altered upon *Lsd1* loss, related to Figure 1.**

(A) Schematic diagram of sgRNAs targeting the regions in the LSD1 protein (left panel) and the exons 1 and 6 of the *Lsd1* genomic sequence. The sgRNAs and PAM sequences are marked in red and blue, respectively. Only *Lsd1* isoform 1 is depicted for simplification.

(B) Agarose gel electrophoresis for validating *Lsd1* KO1 and KO2. PCR products were amplified using the primers P1-P7 (P1, P2, P3, P5, and P6: forward primers; P4 and P7: reverse primers) depicted in panel (A).

(C) Sanger sequencing analysis of *Lsd1* KO1 (top panel) and KO2 (bottom panel) shows insertions and deletions represented in dashes and blue, respectively.

(D) T7 Endonuclease 1 assay of genomic DNA extracted from WT, *Lsd1* KO1 and *Lsd1* KO2 ESCs. Heteroduplexes generated multiple bands after digestion, which are shown in the red arrow. The primers used for PCR amplification were P1 and P4 and P5 and P7.

(E) Representative western blot of LSD1 on whole-cell extracts (WCE) from WT and *Lsd1* KO ESCs. β-ACTIN is used as the loading control.

(F) Immunofluorescence analysis of LSD1 in WT and *Lsd1* KO ESCs. DAPI was used as a nuclear marker. Scale bar, 20μM.

(G) Cell cycle profile in *Lsd1* KO ESCs relative to WT ESCs.

(H and I) Venn diagram showing the overlap of the (H) upregulated and (I) downregulated genes between *Lsd1* KO ESCs (FC > 1.5 and $p < 0.05$).

(J) RT-qPCR analysis of the endoderm markers (*Sox17* and *Gata6*) in WT and *Lsd1* KO ESCs.

Statistical analysis: unpaired t-test (J). ∗$p < 0.05$, ∗∗∗$p < 0.001$, and ∗∗∗∗$p < 0.0001$. Error bars denote mean ± SD; n = 3 (J). Results are one representative of n = 3 independent experiments (E).

**Figure S2.**

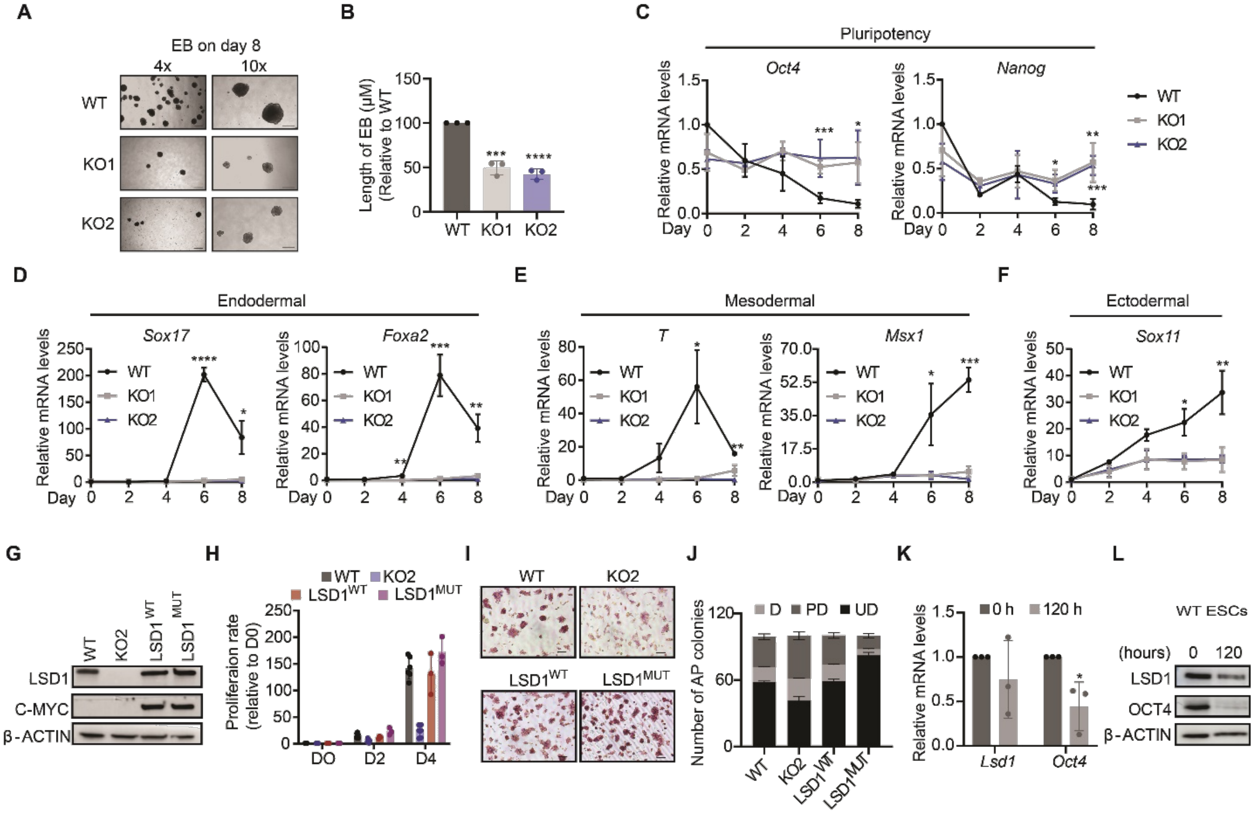


**Figure S2.** ***Lsd1*** **deletion halts differentiation, related to Figure 2.**

(A and B) (A) Representative bright field images at (4x (left) and 10x (right)) magnification and (B) quantification of the size of EB derived from WT and *Lsd1* KO ESCs on the 8th day of differentiation. Scale bars, 200 μM.

(C-F) RT-qPCR analysis of (C) the pluripotency (*Oct4* and *Nanog*), (D) the endodermal (*Sox17* and *Foxa2*), (E) the mesodermal (*T* and *Msx1*), and (F) the ectodermal (*Sox11*) markers in WT, *Lsd1* KO1 and *Lsd1* KO2 ESCs. mRNA levels are relative to the expression of WT at day 0.

(G) Western blot of C-MYC and LSD1 on whole cell extract of WT, *Lsd1* KO2, $LSD1^{WT,}$ and $LSD1^{MUT}$ ESCs. β-ACTIN is as the loading control.

(H) Relative cell proliferation rate of WT, *Lsd1* KO2, $LSD1^{WT,}$ and $LSD1^{MUT}$ ESCs assessed over 4 days.

(I and J) (I) AP staining images and (J) quantification of colonies in WT, *Lsd1* KO2, $LSD1^{WT,}$ and $LSD1^{MUT}$ ESCs. Undifferentiated (UD), partially differentiated (PD), and differentiated (D). Scale bars, 20μM.

(K and L) (K) RT-qPCR of *Lsd1* and (L) Western blot of LSD1 on whole cell extract of gastruloids generated from WT mouse ESCs on indicated time points.

Statistical analysis: unpaired t-test (B-F and K). $*p < 0.05$, $***p < 0.001$, and $****p < 0.0001$. Error bars denote mean ± SD; $n \geq 3$ (B-F and K). Results are one representative of $n = 3$ independent experiments (A, G, I and L).

**Figure S3.**

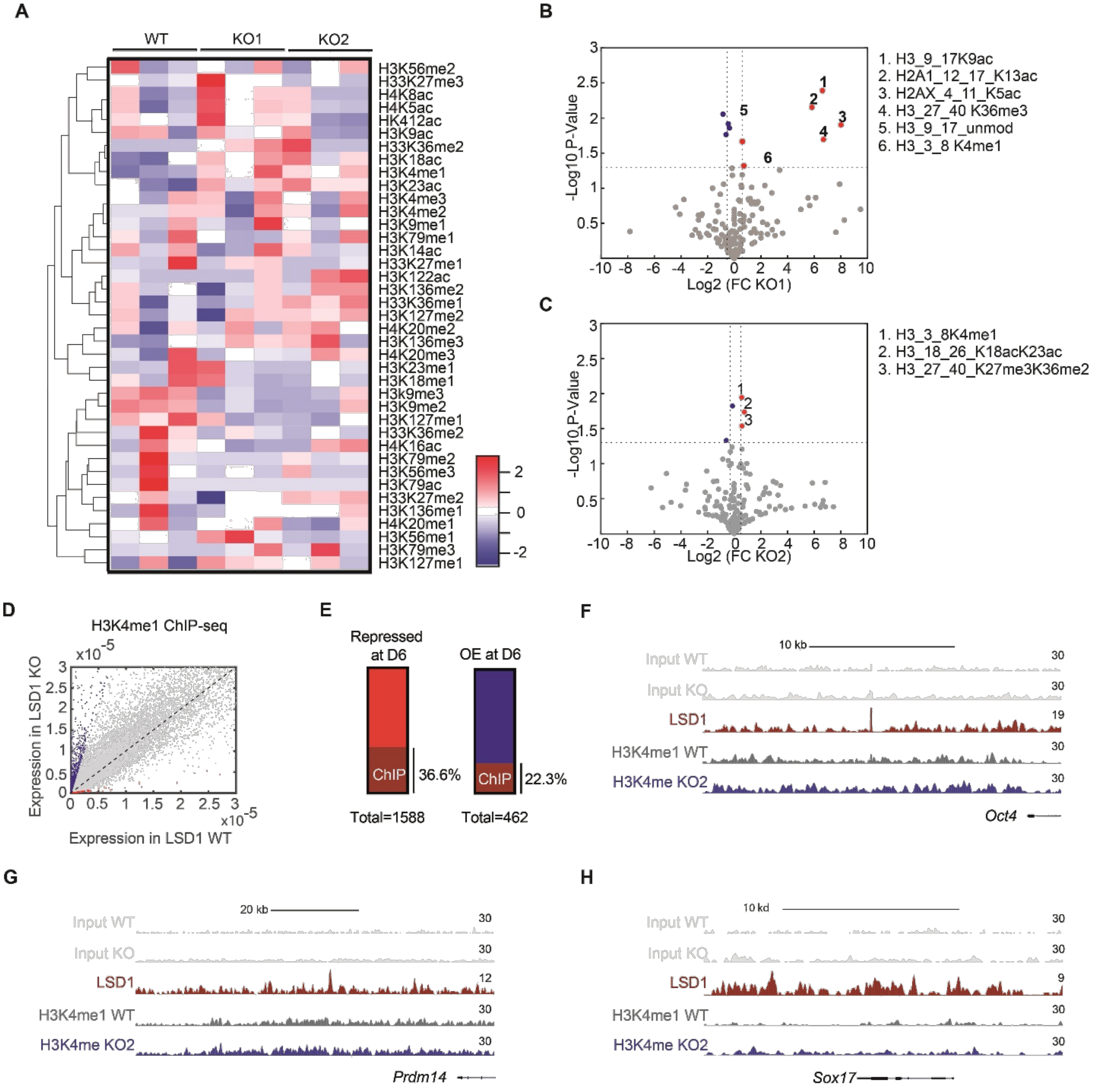


*Figure legends in the next page.*

**Figure S3. Genetic deletion of *Lsd1* affects H3K4me1 and H3K9me2/3, related to Figure 3.**

(A) Heatmap of histone marks in WT, *Lsd1* KO1 and *Lsd1* KO2 ESCs.

(B and C) Volcano plots showing the top enriched histone marks in (B) *Lsd1* KO1 and (C) KO2 relative to WT ESCs. The significant upregulated and downregulated are represented in red and blue respectively ($p < 0.05$ and $FC > 1.5$).

(D) Overlap of *Lsd1* RNA-seq with H3K4me1 ChIP-seq in WT and *Lsd1* KO2 ESCs.

(E) Overlap of LSD1 ChIP-seq with publicly available RNA-seq data on RA-directed differentiation on indicated time points.

(F-H) LSD1 ChIP-seq signal in mouse ESCs and H3K4me1 ChIP-seq signal in WT and *Lsd1* KO2 ESCs at the (F) *Oct4,* (G) *Prdm14 and* (H) *Sox17* enhancers. Respective inputs are depicted in grey.

**Figure S4.**

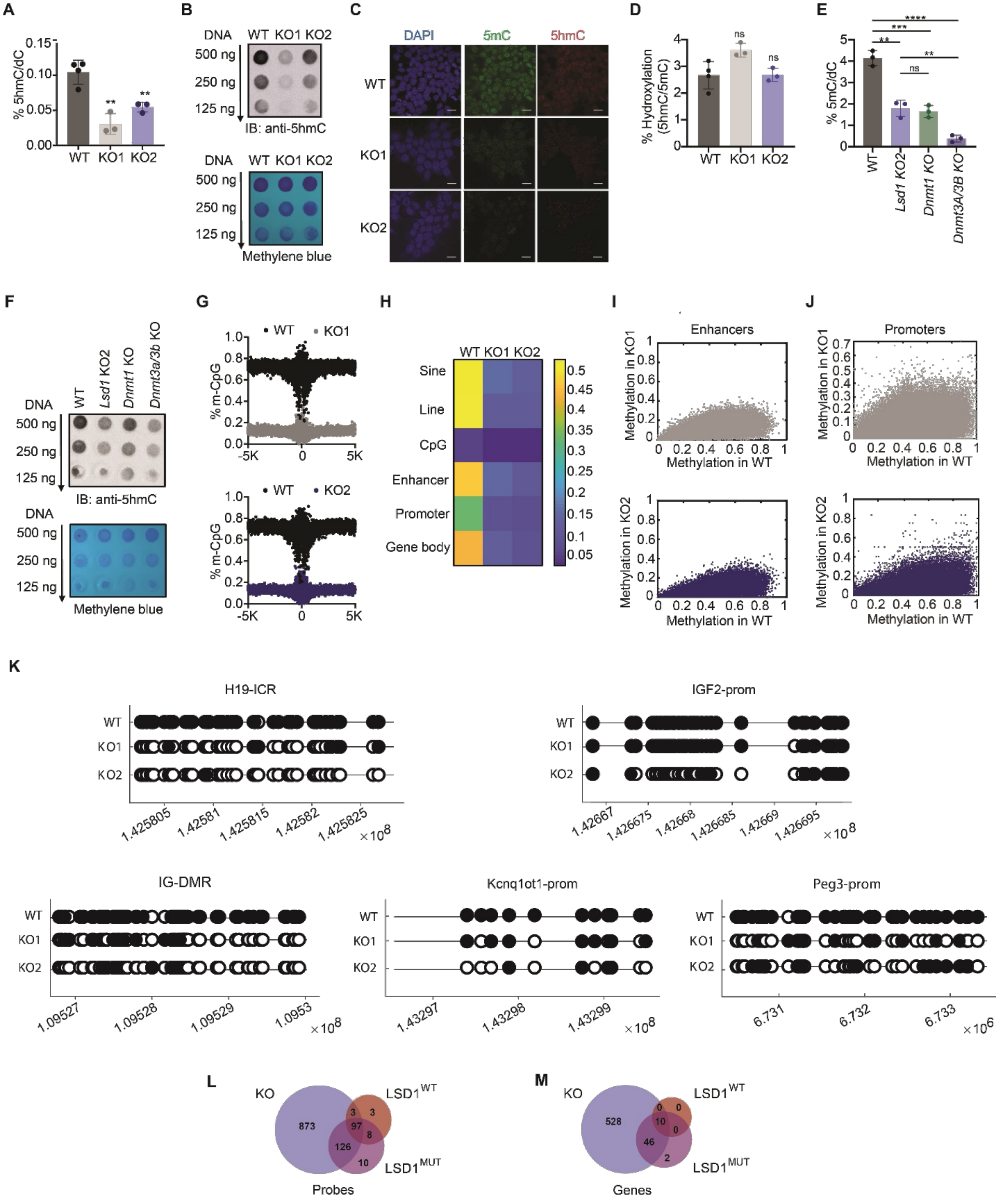


*Figure legends in the next page.*

**Figure S4. Loss of LSD1 affects global DNA methylation, related to Figure 4.**

(A and B) (A) LC-MS/MS quantification (B) DNA dot blot (left panel) of 5hmC on genomic DNA in WT, *Lsd1* KO1 and *Lsd1* KO2 ESCs. 5hmC was normalized against 5mdC. Methylene blue staining was used as the loading control (right panel).

(C) Immunofluorescence of 5mC and 5hmC in WT, *Lsd1* KO1 and *Lsd1* KO2 ESCs. DAPI was used as the nuclear marker. Scale bar, 20μM

(D) Percentage hydroxylation of cysteine in the genomic DNA extracted from WT, *Lsd1* KO1 and *Lsd1* KO2 ESCs.

(E and F) (E) LC-MS/MS quantification and (F) DNA dot blot analysis of 5mC in genomic DNA of WT, *Lsd1* KO2, *Dnmt1* KO, and *Dnmt3a/3b* KO ESCs (top panel). Methylene blue staining was used as the loading control (bottom panel).

(G) Composite plot of methylation in *Lsd1* KO1 (top panel) and *Lsd1* KO2 (bottom panel) compared to WT ESCs across the CpG islands.

(H) Heatmap depicting the DNA methylation distribution in the different regulatory regions in WT, *Lsd1* KO1 and *Lsd1* KO2 ESCs.

(I) Scatter plot of correlation analysis of enhancer methylation in *Lsd1* KO1 (top panel) and *Lsd1* KO2 (bottom panel) compared to WT.

(J) Scatter plot of correlation analysis of promoter methylation in *Lsd1* KO1 (top panel) and *Lsd1* KO2 (bottom panel) compared to WT.

(K) Dot blot of DNA methylation levels in imprinting genes such as *H19*, *Igf2*, *IG-DMR*, *Dlk1-Dio3* and *Peg3.* Methylated and unmethylated CpG are indicated in filled black circles and empty circles.

(L and M) Venn diagram showing overlap of differentially methylated (L) probes (M) genes in *Lsd1* KO2, $LSD1^{WT}$ and $LSD1^{MUT}$ ESCs.

Statistical analysis: unpaired t-test (A, D and E). $**p < 0.01$, $***p < 0.001$, and $****p < 0.0001$. Error bars denote mean ± SD; $n \geq 3$ (A, D and E). Results are one representative of $n = 3$ independent experiments (B, C and F).

**Figure S5.**

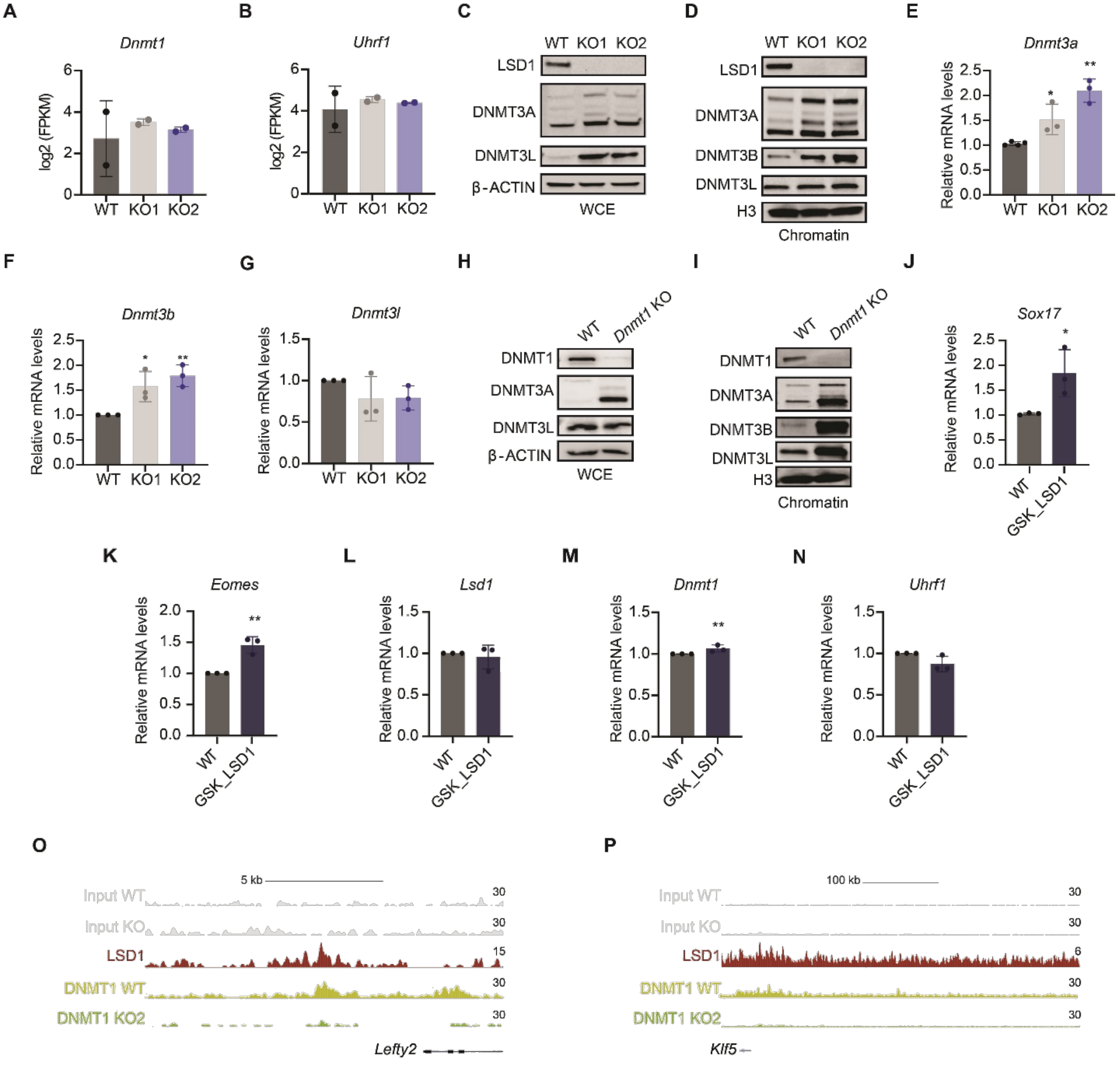


*Figure legends in the next page.*

**Figure S5. LSD1 regulates DNMT1 and UHRF1 protein levels, related to Figure 5.**

(A and B) Expression level (FPKM) of *Dnmt1* and *Uhrf1* in WT, *Lsd1* KO1 and *Lsd1* KO2 ESCs as determined by RNA-seq.

(C and D) Western blots of LSD1, DNMT3A, DNMT3L on the (C) WCE and (D) chromatin fractions of WT and *Lsd1* KOs ESCs. β-ACTIN and H3 are used as the loading controls.

(E-G) RT-qPCR of (E) *Dnmt3a*, (F) *Dnmt3b* and (G) *Dnmt3l* in WT, *Lsd1* KO1 and *Lsd1* KO2 ESCs. mRNA levels are relative to the expression of WT.

(H and I) Western blots of DNMT1, DNMT3A and DNMT3L on the (H) WCE and (I) chromatin fractions of WT and *Lsd1* KO ESCs. β-ACTIN and H3 are used as the loading controls.

(J-N) RT-qPCR of (J) *Sox17,* (K) *Eomes* (L) *Lsd1* (M) *Dnmt1* and (N) *Uhrf1* in WT (vehicle) and inhibitor (GSK_LSD1) treated WT mouse ESCs. mRNA levels are relative to the expression of WT.

(O and P) LSD1 ChIP-seq signal in mouse ESCs and DNMT1 ChIP-seq signal in WT and *Lsd1* KO2 ESCs at the (O) *Lefty2,* and (P) *Klf5* enhancers. Respective inputs are depicted in grey.

Statistical analysis: unpaired t-test (E-G and J-N). *$p<0.05$, **$p < 0.01$. Error bars denote mean ± SD; n =3 (E-G and J-N). Results are one representative of n = 3 independent experiments (C, D, H and I).

**Figure S6.**

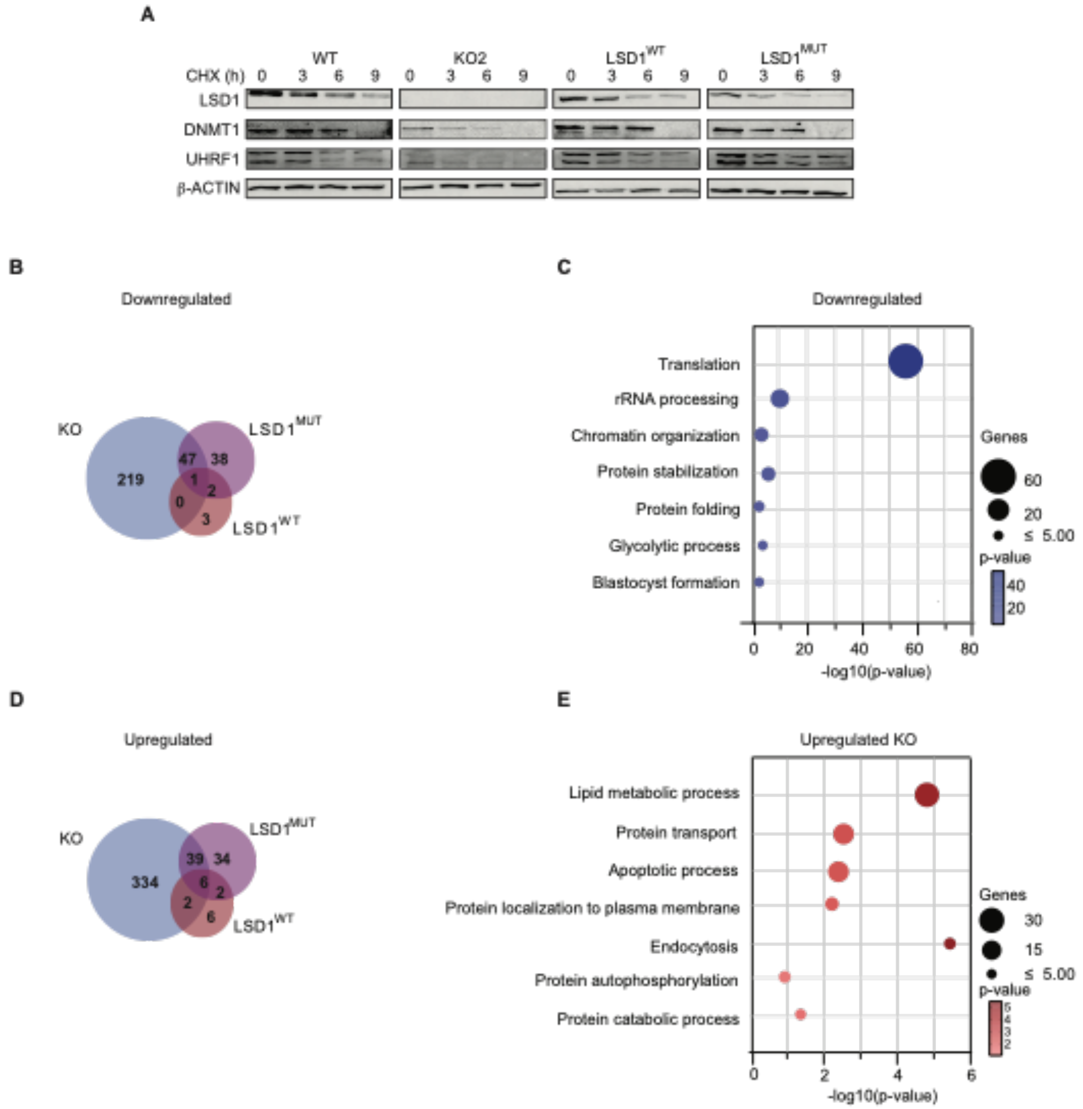


**Figure S6. Silencing of *Lsd1* resulted in decreased DNMT1 and UHRF1 stability, related to Figure 6.**

(A) Western blots of LSD1, DNMT1 and UHRF1 in whole cell extract of WT, *Lsd1* KO2, $LSD1^{WT}$, and $LSD1^{MUT}$ ESCs during a 9 h CHX time course treatment. β-ACTIN is used as the loading control.

(B) Venn diagram showing the overlap of the downregulated proteins in between *Lsd1* KO2, $LSD1^{WT}$, and $LSD1^{MUT}$ ESCs ($p < 0.05$ and FC < 0.8 ).

(C) GO analysis of biological processes related to the downregulated proteins that are exclusive to *Lsd1* KO2 ($p < 0.05$ and FC < 0.8).

(D) Venn diagram showing the overlap of the upregulated proteins in between *Lsd1* KO2, $LSD1^{WT}$, and $LSD1^{MUT}$ ESCs ($p < 0.05$ and FC > 1.2).

(E) GO analysis of biological processes related to the upregulated proteins that are exclusive to *Lsd1* KO2 ($p < 0.05$ and FC > 1.2).

Results are one representative of n = 3 independent experiments (A).
